# Inter-kingdom signaling by the *Legionella* autoinducer LAI-1 involves the antimicrobial guanylate binding protein GBP

**DOI:** 10.1101/2024.09.27.615321

**Authors:** Franziska Solger, Jonas Rauch, Simone Vormittag, Mingzhen Fan, Lyudmil Raykov, Paul Charki, Thierry Soldati, Jürgen Seibel, Hubert Hilbi

## Abstract

The causative agent of Legionnaires’ disease, *Legionella pneumophila*, is an amoebae-resistant environmental bacterium, which replicates intracellularly in a distinct compartment, the “*Legionella*-containing vacuole” (LCV). *L. pneumophila* employs the α-hydroxyketone compound LAI-1 (*Legionella* autoinducer-1) for intra-species and inter-kingdom signaling. LAI-1 promotes intracellular replication and inhibits the migration of mammalian cells and *Dictyostelium discoideum*. In this study, we revealed that LAI-1 and “clickable” azido-LAI-1 derivatives inhibit the migration of *D. discoideum* and localize to LCVs. Azido-LAI-1 colocalizes with the LCV markers calnexin, P4C, and AmtA, but not with mitochondrial or lipid droplet markers. Intriguingly, LAI-1 dependent inhibition of *D. discoideum* migration involves the single guanylate-binding protein (GBP), a member of the GBP family of large GTPases, which in metazoan organisms promote cell autonomous immunity. *D. discoideum* lacking GBP (Δ*gnbp*) allows more efficient intracellular replication of *L. pneumophila*, without apparently compromising LCV remodeling or integrity, and GBP-GFP localizes to the ER at LCV-ER membrane contact sites (MCS). However, the peri-LCV localization of LAI-1 and GBP is not mutually dependent. Synthetic LAI-1 inhibits the expansion/remodeling of LCVs (but not vacuoles harboring avirulent *L. pneumophila*) in a GBP-dependent manner. Taken together, the work shows that LAI-1 localizes to LCVs, and LAI-1-dependent inter-kingdom signaling involves *D. discoideum* GBP, which localizes to LCV-ER MCS and acts as an antimicrobial factor by restricting the intracellular growth of *L. pneumophila*.

**Author Summary:** Small molecule inter-kingdom signaling between pathogens and host cells represents a crucial but only partly understood aspect of microbial virulence. The amoeba-resistant opportunistic pathogen *Legionella pneumophila* employs the compound LAI-1 (*Legionella* autoinducer-1) for intra-species and inter-kingdom signaling. In metazoan cells, the conserved and wide-spread family of guanylate-binding protein (GBP) large GTPases usually comprises several distinct paralogues, which are implicated in pathogen detection, inflammation, cell death pathways, and cell autonomous immunity. In the social amoeba *Dictyostelium discoideum*, only a single *GBP* gene of unknown function is present. Using approaches from organic chemistry, genetics, cell biology and infection biology, we reveal that GBP is involved in the inhibition of *D. discoideum* migration and pathogen vacuole expansion/remodeling by LAI-1 as well as in intracellular growth of *L. pneumophila*. This study provides a novel link between small molecule inter-kingdom signaling and GBP-dependent cell autonomous immunity.

## Introduction

Small molecule inter-kingdom communication between bacterial pathogens and eukaryotic target cells represents a crucial aspect of microbial pathogenesis (Hochstrasser & Hilbi, 2017; Kendall & Sperandio, 2016; Ng & Bassler, 2009; Pacheco & Sperandio, 2009; Shank & Kolter, 2009). Zoonotic enteropathogenic bacteria such as *Salmonella enterica* or pathogenic *Escherichia coli*, as well as environmental bacteria such as *Vibrio cholerae* or *Legionella pneumophila* employ small molecule inter-kingdom signaling as a virulence strategy.

In the environment, the opportunistic pathogen *L. pneumophila* replicates in free-living protozoa, including the amoebae *Acanthamoeba castellanii* and *Dictyostelium discoideum* (Boamah *et al*, 2017; Hoffmann *et al*, 2014b; Swart *et al*, 2018). Upon inhalation of *L. pneumophila*-contaminated aerosols, the bacteria grow within and destroy lung macrophages, thereby causing a severe pneumonia termed Legionnaires’ disease (Hilbi *et al*, 2011; Mondino *et al*, 2020; Newton *et al*, 2010). To govern the interactions with these evolutionarily distant phagocytes, amoebae and macrophages, *L. pneumophila* employs the Icm/Dot type IV secretion system (T4SS), which translocates more than 300 “effector proteins” into the host cells (Hilbi & Buchrieser, 2022; Lockwood *et al*, 2022; Personnic *et al*, 2016; Qiu & Luo, 2017; Swart *et al*, 2020). Some of these effectors have been described to target crucial host processes and to promote the formation of a replication-permissive compartment termed the *Legionella*-containing vacuole (LCV) (Asrat *et al*, 2014; Hubber & Roy, 2010; Steiner *et al*, 2018; Swart & Hilbi, 2020). Critical steps in the LCV formation are the modulation of the small GTPases Arf1 and Rab1 (Goody & Itzen, 2013), phosphoinositide conversion from phosphatidylinositol 3-phosphate (PdtIns(3)*P*) to PdtIns(4)*P* (Weber *et al*, 2018; Weber *et al*, 2014; Weber *et al*, 2006), the formation of membrane contact sites (MCS) with the endoplasmic reticulum (ER) (Vormittag *et al*, 2023a; Vormittag *et al*, 2023b), and intimate interactions with lipid droplets (LD) (Hüsler *et al*, 2023a; Hüsler *et al*, 2023b; Hüsler *et al*, 2021).

*L. pneumophila* employs the *Legionella* quorum sensing (Lqs) system for small molecule signaling, which produces, detects and responds to an organic α-hydroxyketone compound called *Legionella* autoinducer-1 (LAI-1, 3-hydroxypentadecane-4-one) (Hochstrasser & Hilbi, 2017; Michaelis *et al*, 2024b; Tiaden & Hilbi, 2012; Tiaden *et al*, 2010a). LAI-1 is secreted and delivered to prokaryotic and eukaryotic cells by bacterial outer membrane vesicles (Fan *et al*, 2023). The Lqs system comprises the autoinducer synthase LqsA (Spirig *et al*, 2008), the homologous membrane-bound sensor histidine kinases LqsS (Tiaden *et al*, 2010b) and LqsT (Kessler *et al*, 2013), and the cognate cytosolic response regulator LqsR (Tiaden *et al*, 2008; Tiaden *et al*, 2007), which dimerizes upon phosphorylation and harbors an output domain resembling nucleotide-binding domains (Hochstrasser *et al*, 2020; Schell *et al*, 2014).

The Lqs system is linked through the pleiotropic transcription factor LvbR to signaling pathways involving the inorganic gas nitric oxide (NO) (Hochstrasser *et al*, 2019). These signaling pathways comprise three distinct NO receptors upstream of two-component systems converging on c-di-GMP metabolism (Michaelis *et al*, 2024a). NO signaling regulates virulence, motility, biofilm formation and dispersal, as well as phenotypic heterogeneity of *L. pneumophila*. Taken together, LAI-1-dependent quorum sensing is linked to NO and c-di-GMP signaling jointly regulating a plethora of *L. pneumophila* traits (Hochstrasser & Hilbi, 2020; Michaelis *et al*., 2024b; Striednig & Hilbi, 2022).

LAI-1 and the Lqs-LvbR signaling network promote intra-species communication and regulate various features of *L. pneumophila*, including virulence (Hochstrasser & Hilbi, 2017; Personnic *et al*, 2018), motility and flagellum production (Schell *et al*, 2016), phase switch and temperature-dependent cell density (Hochstrasser & Hilbi, 2022; Tiaden *et al*., 2007), expression of a 133 kb genomic “fitness island” and natural competence for DNA uptake (Hochstrasser & Hilbi, 2017; Personnic *et al*., 2018). Moreover, the Lqs-LvbR network also regulates phenotypic heterogeneity and the occurrence of functionally different *L. pneumophila* subpopulations (e.g., “persisters”) upon infection of amoebae and macrophages (Personnic *et al*, 2019; Striednig *et al*, 2021) as well as in biofilms and under sessile conditions (Personnic *et al*, 2021). Finally, LAI-1 and the Lqs-LvbR signaling network govern inter-kingdom communication by modulating the motility of eukaryotic cells, including *D. discoideum*, macrophages, or epithelial cells (Simon *et al*, 2015), intracellular replication (Fan *et al*., 2023), and migration of *A. castellanii* through *L. pneumophila* biofilms (Hochstrasser *et al*, 2022). Specifically, LAI-1 inhibits epithelial cell migration through a pathway requiring the scaffold protein IQGAP1, the small GTPase Cdc42 (but not RhoA or Rac1), as well as the Cdc42-specific guanine nucleotide exchange factor (GEF) ARHGEF9 (**Fig. 1A**).

**Fig. 1.**
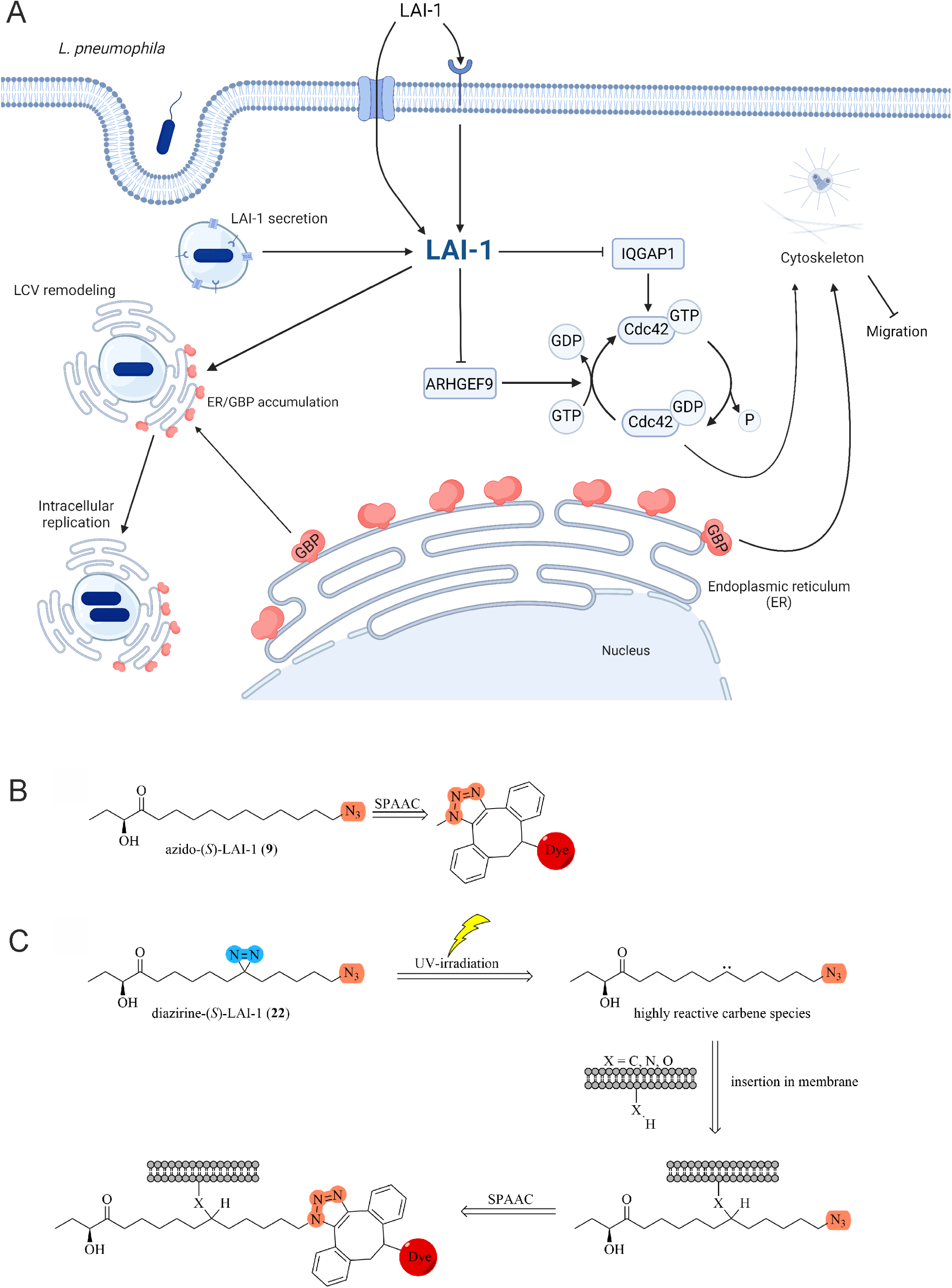
LAI-1-dependent inter-kingdom signaling and role of GBP. (**A**) LAI-1-dependent inter-kingdom signaling of *L. pneumophila* comprises cell migration inhibition and cytoskeleton disintegration. LAI-1 is detected and/or taken up by eukaryotic host cells by unknown mechanisms. The single *D. discoideum* large GTPase guanylate-binding protein, DdGPB, restricts intracellular growth of *L. pneumophila*, localizes to LCV-ER membrane contact sites and is required for LAI-1-dependent LCV size remodeling. Thus, LAI-1 links small molecule inter-kingdom signaling and GBP-dependent cell autonomous immunity. (**B**) Azido-(*S*)-LAI-1 can be attached to various conjugation partners (e.g., dyes) using bioorthogonal strain-promoted alkyne-azide cycloadditions (SPAAC). (**C**) The diazirine function of azido-diazirine-LAI-1 is stimulated by UV light and forms a carbene by releasing nitrogen. The highly reactive carbene can interact with various chemical moieties and thus covalently binds to its biological environment. The covalently fixed azido-LAI-1-derivative can then be attached to various conjugation partners (e.g., dyes, biotin) using SPAAC.

To counteract microbial assaults, eukaryotic cells have evolved a sophisticated array of cell-autonomous defense mechanisms. Guanylate-binding protein (GBP) large GTPases comprise a conserved and wide-spread family of antimicrobial nanomachines (Kutsch & Coers, 2021; Tretina *et al*, 2019). In mammalian cells, several paralogues of interferon-γ (IFN-γ)-induced GBPs are usually present, e.g., 7 human and 11 mouse GBPs have been identified (Kirkby *et al*, 2023). Increasing evidence suggests that these GBPs represent cytosolic pattern recognition receptors (PRRs), which bind conserved microbial structures like lipopolysaccharide (LPS) of Gram-negative bacteria and directly kill microorganisms (Kutsch *et al*, 2020; Santos *et al*, 2020). Furthermore, GBPs also disrupt pathogen vacuoles and/or activate the inflammasome and pyroptotic cell death to protect eukaryotic cells from invading microorganisms (Rafeld *et al*, 2021; Tretina *et al*., 2019). In human macrophages, GBP1 is required for IFN-γ-induced inflammasome responses against *L. pneumophila* and co-localizes with LCVs to promote the rupture of the pathogen vacuole (Bass *et al*, 2023). In mouse macrophages, GBPs are also required for inflammasome-dependent clearance of *L. pneumophila* but dispensable for LCV rupture (Liu *et al*, 2018). GBPs comprise a globular N-terminal GTPase domain followed by an extended C-terminal α-helical domain (Kirkby *et al*., 2023; Tretina *et al*., 2019). Upon GTP binding, GBPs form homodimers, which further assemble into large, multimeric complexes on microbial or host membranes. However, the target specificity and mode of action of GBPs are incompletely understood at present.

The *D. discoideum* genome encodes a single GBP homologue termed DdGBP (Dunn *et al*, 2017; Katic *et al*, 2021), which is most closely related to human GBP3 (26% protein identity, *e*-value 3 × 10^-38^). The corresponding *gnbp* gene (Q54TN9 / DDB_G0281639), encodes a protein of 796 amino acids (92,707 Da). DdGBP is predicted to harbor a C-terminal signal peptide with a basic double lysine motif and an N-terminal transmembrane domain; yet, the protein/gene has not been characterized. Intriguingly, however, GBP was identified by mass spectrometry in the proteome of purified macropinosomes (Journet *et al*, 2012) and of purified LCVs isolated from *L. pneumophila*-infected *D. discoideum* (Hoffmann *et al*, 2014a). In this study, we reveal that (i) LAI-1 and GBP-GFP localize to LCVs, (ii) GBP restricts intracellular growth of *L. pneumophila*, and (iii) LAI-1-dependent inhibition of *D. discoideum* migration and LCV size expansion/remodeling involves GBP. Hence, LAI-1-dependent inter-kingdom signaling involves the single antimicrobial *D. discoideum* GBP.

## Results

### LAI-1 and azido-derivatives inhibit amoeba migration and localize to LCVs

Synthetic LAI-1 inhibits the migration of *D. discoideum* in a dose-dependent manner (Simon *et al*., 2015). To further characterize LAI-1-dependent interkingdom signaling, we sought to introduce functional LAI-1 derivatives, “clickable” azido-LAI-1 and UV-activatable azido-diazirine-LAI-1 (termed diazirine-LAI-1 in this study; see Materials and Methods section for synthesis details). Previously, we have shown that functionalized clickable azido-sphingolipids are incorporated into cellular membranes similar to control sphingolipids without modification allowing the visualization of sphingomyelin distribution and sphingomyelinase activity in infection processes (Fink *et al*, 2021; Rühling *et al*, 2024; Sternstein *et al*, 2023).

LAI-1 was synthetically equipped with an azide group, which enables the spontaneous click reaction between the azide and dibenzocyclooctyne (DBCO) dyes (**Fig. 1B**). The terminal azide promotes minimally invasive bioorthogonal strain-promoted alkyne-azide cycloadditions (SPAAC), allowing the conjugation with fluorescent dyes (e.g., DIBO594). Since conjugation with large molecules has a major influence on the substance properties and biological targets might diffuse in the cell, covalent photo-crosslinking with bifunctional LAI-1 is desirable. Accordingly, we further synthesized UV-activatable azido-diazirine-LAI-1, allowing the spontaneous cross-linking of LAI-1 with membranes and proteins by photo-activation (**Fig. 1C**). Diazirine-LAI-1 can be excited by UV radiation, whereupon a highly reactive carbene is formed, which crosslinks with functional groups in close proximity. This covalent modification prevents diffusion from the target, and the additional azide group enables coupling with various conjugation partners via SPAAC.

Treatment of *D. discoideum* Ax3 with 10 µM LAI-1, azido-LAI-1, or diazirine-LAI-1 inhibited the migration of single amoebae (**Fig. 2A**). Compared to the solvent control, the velocity of *D. discoideum* treated with LAI-1, azido-LAI-1, or diazirine-LAI-1 was significantly reduced by ca. 30% (**Fig. 2B**). LAI-1 and the derivatives reduced amoebae velocity to a similar extent, indicating that the inter-kingdom signaling activity of the derivatives is not compromised. Analogously, azido-LAI-1 triggered the luminescence of a *Vibrio cholerae* α-hydroxyketone reporter strain in a concentration-dependent manner (**Fig. S1A**), similarly to what has been previously found for LAI-1 (Fan *et al*., 2023). Diazirine-LAI-1 triggered luminescence of the *V. cholerae* reporter strain less effectively but still above background level. In summary, azido- and diazirine-azido-LAI-1 derivatives are biologically active in inter-kingdom as well as inter-bacterial signaling, and hence, the derivatives are valid tools to study the effects of LAI-1 on eukaryotic and prokaryotic cells.

**Fig. 2.**
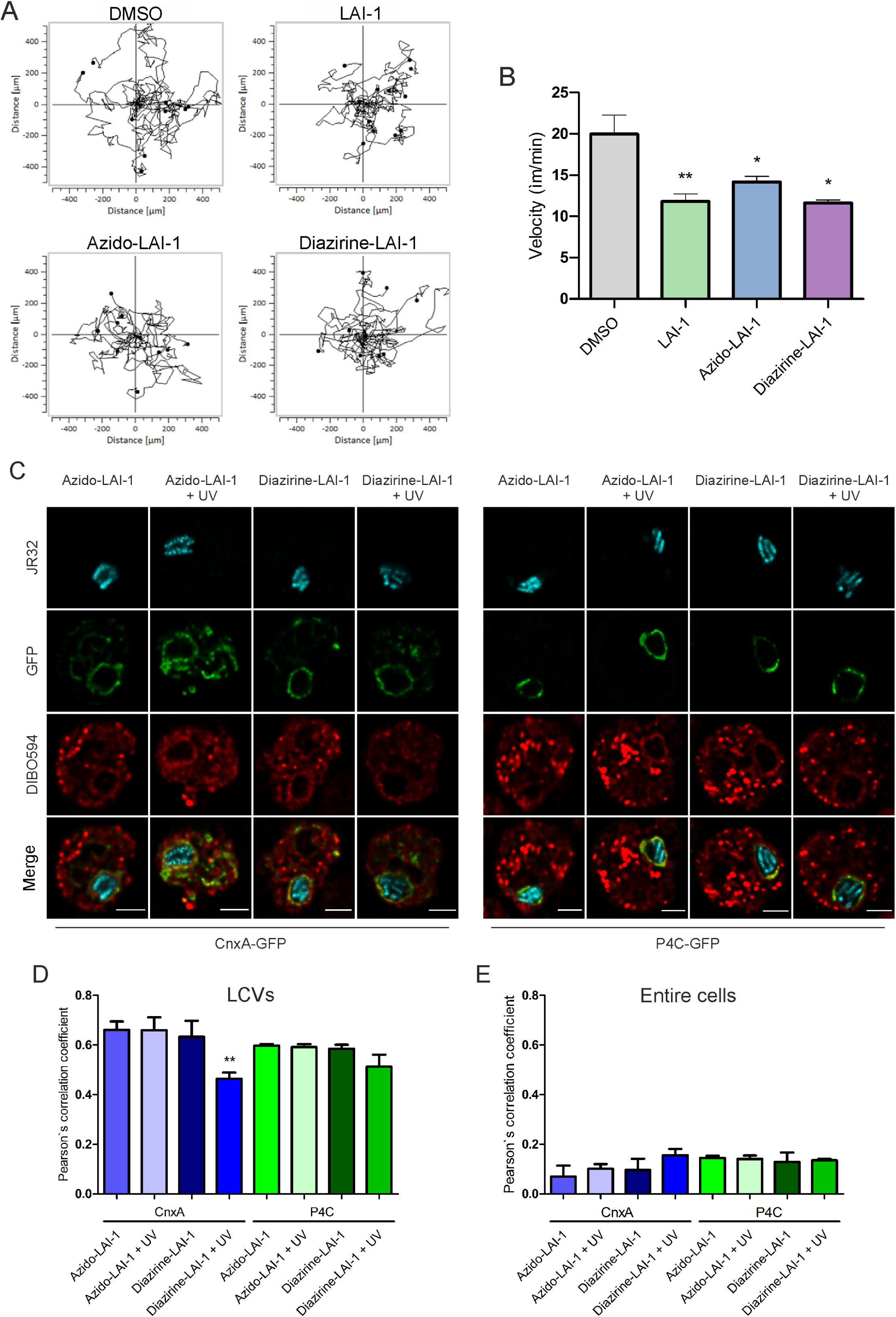
LAI-1 and azido-derivatives inhibit amoeba migration and localize to LCVs. **(A)** *D. discoideum* Ax3 producing GFP (pSW102) was treated (10 µM, 1 h) with (*S*)-LAI-1, (*S*)-azido-LAI-1 (termed “azido-LAI-1)”, (*S*)-azido-diazirine-LAI-1 (termed “diazirine-LAI-1”) or DMSO (solvent control), and single cell migration was recorded continuously for 2 h with 2 min time interval. (**B**) Amoeba velocity was quantified using the ImageJ manual tracker and Ibidi chemotaxis software. (**C**) *D. discoideum* Ax2 producing calnexin (CnxA)-GFP (pAW016) or P4C-GFP (pWS034) was treated with azido-LAI-1 or diazirine-LAI-1 (10 µM, 1 h), infected (MOI 5, 8 h) with mCerulean-producing *L. pneumophila* JR32 (pNP99), exposed to UV light (5 min), clicked with DIBO594 dye, fixed (24 h p.i.) and analyzed by confocal laser scanning microscopy. Scale bars, 3 μm. The colocalization of azido-LAI-1 (DIBO594 dye) or diazirine-LAI-1 (DIBO594 dye) with CnxA-GFP or P4C-GFP was quantified by Pearson’s correlation coefficient for (**D**) LCVs or (**E**) entire cells. Data shown are (A) representative of at least three biological replicates, or (B, D, E) means and standard deviations of biological triplicates (Student’s t-test; *, *p* ≤ 0.05; **, *p* ≤ 0.01).

The clickable LAI-1 derivatives were then used to assess the intracellular localization of LAI-1. Upon addition of clickable DIBO594 dye to *D. discoideum* pre-treated with 10 µM azido-LAI-1 or LAI-1 as a negative control, the clickable LAI-1 derivative but not the LAI-1 control labeled the amoebae (**Fig. S1B**). To test the intracellular localization of LAI-1, *D. discoideum* amoebae were treated with 10 µM azido-LAI-1 or diazirine-LAI-1 and infected with mCerulean-producing *L. pneumophila* JR32. At 8 h post infection (p.i.), some of the infected amoebae were UV-irradiated, further incubated for 24 h (with clickable DIBO594 dye for the last 30 min), fixed, and imaged by confocal microscopy (**Fig. 2C**). This approach revealed that LAI-1 localizes to LCVs as judged from the co-localization with LCV-associated ER (calnexin/CnxA) as well as with the PtdIns(4)*P*-positive LCV membrane (P4C) (**Fig. 2D**). In addition to LCVs, LAI-1 also localizes to other membrane-bound cellular compartments throughout the cells (**Fig. 2E**). Upon treatment of the samples with UV, diazirine-LAI-1, but not azido-LAI-1, co-localized less extensively with LCV-associated calnexin and PtdIns(4)*P* (**Fig. 2D**), validating that UV specifically affects diazirine-but not azido-LAI-1. Moreover, the results are in agreement with the notion that at later stages of the infection, the LAI-1-binding LCV membranes dynamically reorganize. In summary, LAI-1 and clickable azido-derivatives inhibit *D. discoideum* migration and localize to LCVs in the amoebae.

### LAI-1 localizes to LCV-ER membrane contact sites and the LCV membrane

Next, we sought to assess the subcellular localization of LAI-1 in more detail and at different early timepoints p.i. To this end, we used *D. discoideum* producing calnexin-GFP (ER), P4C-GFP (LCV membrane/PtdIns(4)*P*), GREMIT (mitochondria), or Plin-GFP (lipid droplets/LD). The amoebae were treated with 10 µM azido-LAI-1, infected with mCerulean-producing *L. pneumophila* JR32 and clicked with DIBO594 dye (**Fig. 3A**). Fluorescence intensity profiles and the quantification of co-localization by Pearson’s correlation coefficient revealed that in *L. pneumophila*-infected *D. discoideum*, LAI-1 co-localizes with calnexin-GFP as well as with P4C-GFP on LCVs (**Fig. 3B**) and in whole cells (**Fig. 3C**). Throughout the infection (0.5-8 h p.i.), LAI-1 did not co-localize with either GREMIT or Plin-GFP, indicating that LAI-1 does not localize to mitochondria or LD. Interestingly, LAI-1 specifically localizes to LCVs harboring wild-type *L. pneumophila* but not to the AmtA-positive vacuole harboring Δ*icmT* mutant bacteria lacking a functional T4SS (**Fig. 3B**). In summary, these results reveal that LAI-1 co-localizes with the ER at LCV-ER MCS as well as with the PtdIns(4)*P*-positive LCV membrane.

**Fig. 3.**
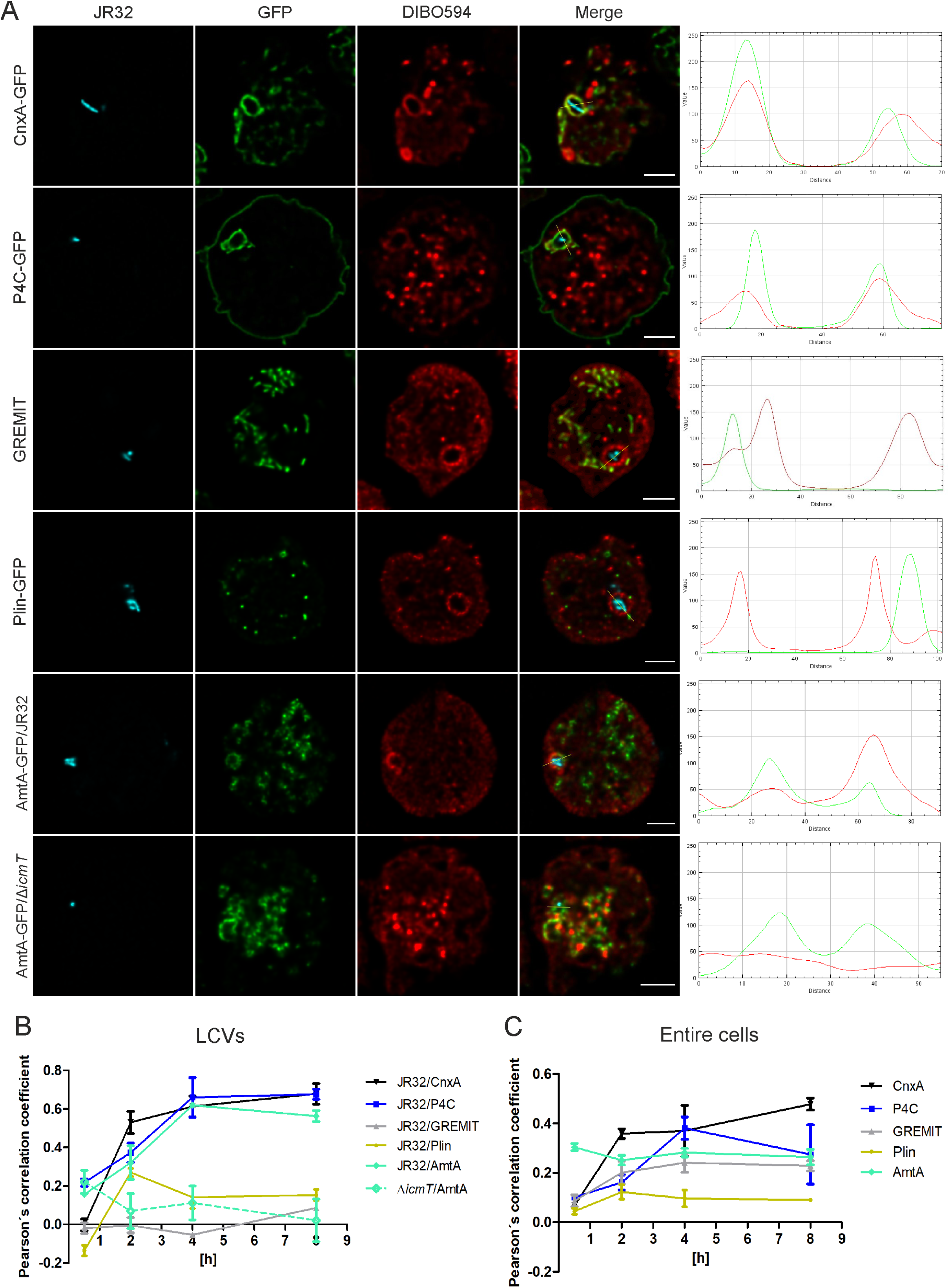
LAI-1 localizes to LCV-ER membrane contact sites and the LCV membrane. **(A)** *D. discoideum* Ax2 producing calnexin (CnxA)-GFP (ER; pAW016), P4C-GFP (LCV membrane; pWS034), GREMIT (mitochondria), GFP-Plin (LD; pHK101), or AmtA (early endosomes; pDM1044-AmtA-mCherry) was treated with clickable azido-LAI-1 (10 µM, 1 h), infected (MOI 5, 4 h) with mCerulean-producing *L. pneumophila* JR32 or Δ*icmT* (pNP99), clicked with DIBO594 dye, fixed, and analyzed by confocal microscopy. Scale bars, 3 µm (fluorescence images; left panels). Fluorescence intensity profiles were generated for the GFP fusion proteins and DIBO594 dye using the RGB profile from ImageJ (right panels). The co-localization of azido-LAI-1 (DIBO594 dye) with different organelle markers was quantified by Pearson’s correlation coefficient for (**B**) LCVs or (**C**) entire cells (0.5-8 h p.i.). Data shown are means and standard deviations of biological triplicates.

### LAI-1-dependent inhibition of *Dictyostelium* migration involves GBP

Searching for host factors that might affect the intracellular growth of *L pneumophila* in *D. discoideum* and/or the effects of LAI-1 on the amoebae, we constructed and tested a *D. discoideum* mutant strain lacking the single GBP family large GTPase. The *gnbp* gene was deleted from the *D. discoideum* Ax2 genome by double homologous recombination, yielding the Δ*gnbp* mutant strain. The mutant strain lacking GBP grew like the parental amoebae under axenic conditions. Compared to the parental strain, Δ*gnbp* amoeba showed increased random migration (**Fig. 4A**) and moved with significantly increased velocity (**Fig. 4B**). The Δ*gnbp* migration phenotype was reverted by plasmid-borne production of GBP-GFP, validating the genetic setup of the mutant strain and indicating that the C-terminal fusion of GBP with GFP is functional. Taken together, DdGBP negatively regulates random amoeba motility.

**Fig. 4.**
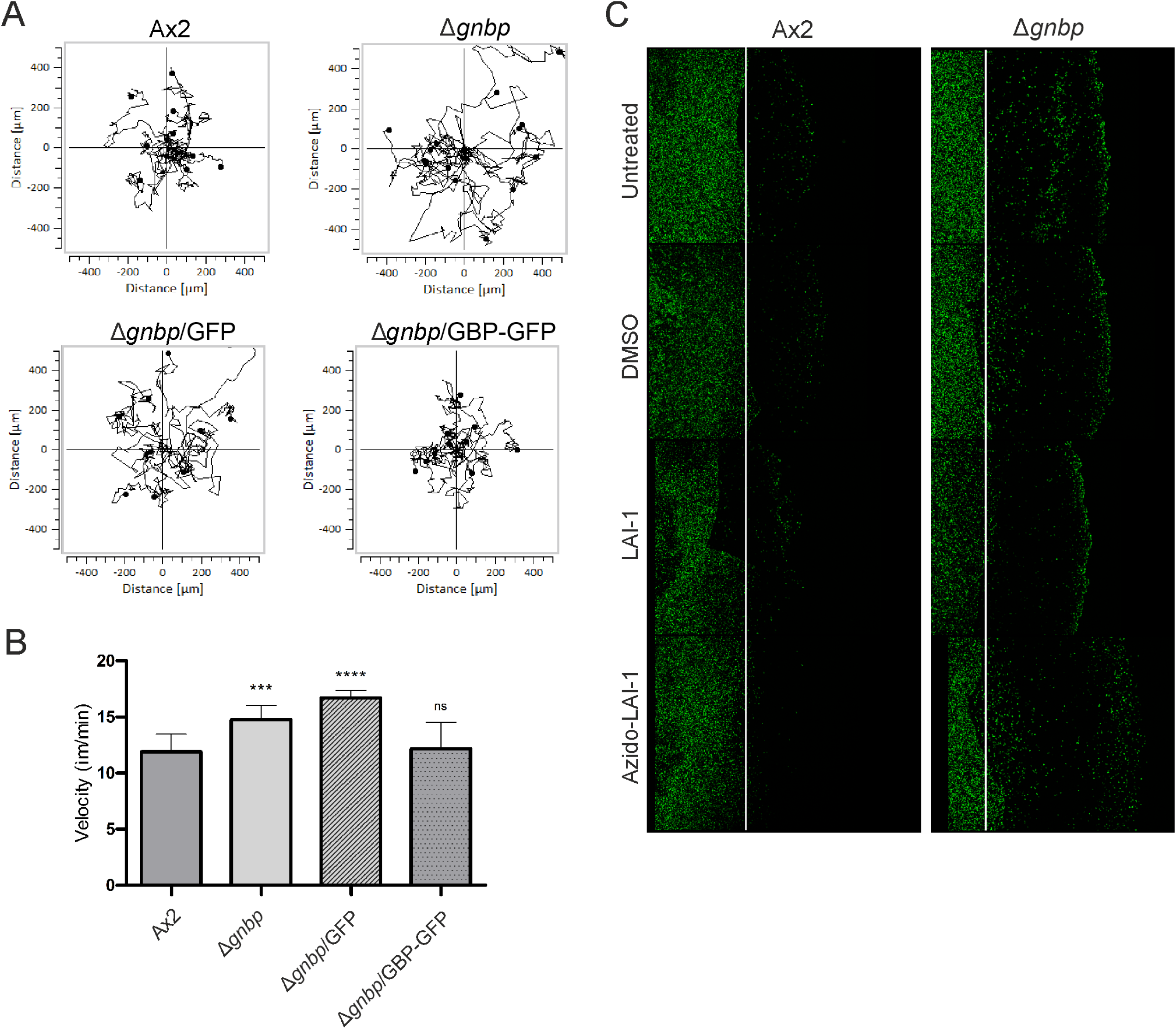
LAI-1-dependent inhibition of *Dictyostelium* migration involves GBP. (**A**) Single cell migration of *D. discoideum* Ax2, Δ*gnbp* or Δ*gnbp* producing GFP (pDM317) or GBP-GFP was recorded continuously for 2 h with 2 min time interval. (**B**) Amoeba velocity was quantified using the ImageJ manual tracker and Ibidi chemotaxis software. (**C**) *D. discoideum* Ax2 or Δ*gnbp* producing GFP (pDM317) was left untreated or treated (10 µM, 1 h) with LAI-1, azido-LAI-1, or DMSO (solvent control), and cell migration towards 1 mM folate was assessed by under-agarose assay (4 h p.i.). The white lines represent the edge of the sample wells. Data shown are (A) representative of at least three biological replicates, or (B) means and standard deviations of biological triplicates (Student’s t-test; ***, *p* ≤ 0.001; ****, *p* ≤ 0.0001).

Next, we assessed the migration towards folate of *D. discoideum* Ax2 and Δ*gnbp* amoebae left untreated or treated with LAI-1 or azido-LAI-1 (**Fig. 4C**). Similarly to random migration, amoebae lacking GBP showed enhanced chemotactic migration towards folate. Interestingly, compared to the parental amoebae, Δ*gnbp* cells showed a reduced response to treatment with LAI-1 or azido-LAI-1, revealing the involvement of GBP in LAI-1-dependent migration inhibition. While the chemotactic migration of the parental *D. discoideum* strain was dose-dependently impaired by 1-10 µM LAI-1, the migration of the Δ*gnbp* strain was not inhibited (**Fig. S2**). In summary, these results indicate that the single *D. discoideum* GBP is implicated in random and chemotactic amoebae migration as well as in LAI-1-dependent inhibition of migration.

### GBP restricts growth of *L. pneumophila* and co-localizes with LAI-1 and the ER at LCV-ER MCS

Given the role of GBP for *D. discoideum* migration and LAI-1-dependent migration inhibition, we next assessed whether GBP affects the intracellular growth of *L. pneumophila* (**Fig. 5A**). Compared to the parental *D. discoideum* strain, *L. pneumophila* grew significantly more efficiently in the Δ*gnbp* mutant strain, indicating that GBP restricts intracellular growth of the pathogen. In contrast, the avirulent *L. pneumophila* Δ*icmT* mutant strain did not grow in the mutant amoebae (**Fig. S3A**). *L. pneumophila* was taken up with the same efficiency by the parental *D. discoideum* strain and Δ*gnbp* mutant amoebae (**Fig. S3BC**), and therefore, the enhanced intracellular bacterial growth is not due enhanced uptake. Treatment of *D. discoideum* Ax2 or Δ*gnbp* with LAI-1 prior to an infection did neither substantially alter the course of the infection (**Fig. 5A**) nor bacterial uptake (**Fig. S3BC**). Intriguingly, while DdGBP restricted the intracellular growth of *L. pneumophila*, it did not affect the intracellular growth of the amoeba-resistant pathogen *Mycobacterium marinum* (**Fig. S4A**). In summary, *L. pneumophila* wild-type but not Δ*icmT* mutant bacteria grow more efficiently in the Δ*gnbp* mutant strain, while bacterial uptake was not affected. These findings identify DdGBP as an antimicrobial factor.

**Fig. 5.**
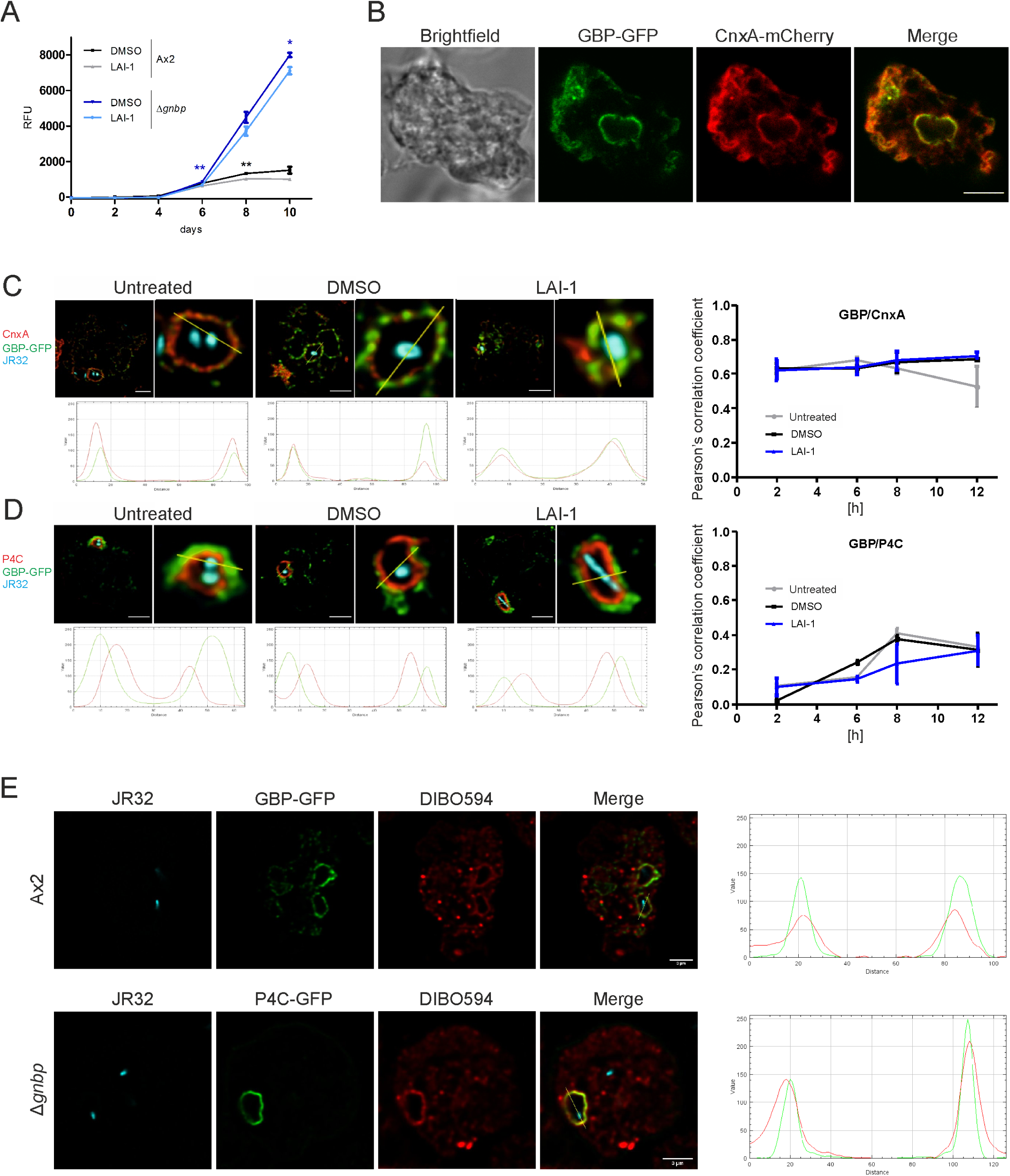
GBP restricts intracellular growth of *L. pneumophila* and co-localizes with LAI-1 and the ER at LCV-ER MCS. (**A**) *D. discoideum* Ax2 or Δ*gnbp* was treated with LAI-1 (10 µM, 1 h) or DMSO (solvent control), infected (MOI 1, 10 d) with GFP-producing *L. pneumophila* JR32 (pNT28), and intracellular replication was assessed by RFU. Data shown are means and standard deviations of biological triplicates (Student’s t-test; *, *p* ≤ 0.05; **, *p* ≤ 0.01). (**B**) Dually labeled *D. discoideum* Ax2 producing GBP-GFP and calnexin (CnxA)-mCherry (pAW012) was fixed and analyzed by confocal microscopy. Scale bar, 5 µm. Dually labeled *D. discoideum* Ax2 producing GBP-GFP and (**<u>C</u>**) CnxA-mCherry (pAW012) or (**D**) P4C-mCherry (pWS032) was left untreated or treated with LAI-1 (10 µM, 1 h) or DMSO (solvent control), infected (MOI 5, 2-12 h) with mCerulean-producing *L. pneumophila* JR32 (pNP99), fixed, and analyzed by confocal microscopy (8 h p.i., left panels). Scale bars, 5 µm. The colocalization of GBP with the ER (CnxA; C) or with the LCV membrane (P4C; D) was quantified by Pearson’s correlation coefficient (right panels). Fluorescence intensity profiles were generated for the GFP fusion proteins and CnxA (C; lower panels) or P4C (D; lower panels) using the RGB profile from ImageJ. Data shown (right panels) are biological triplicates of means and standard deviations. (**E**) *D. discoideum* Ax2 or Δ*gnbp* producing GBP-GFP or P4C-GFP, respectively, was treated with clickable azido-LAI-1 (10 µM, 1 h), infected (MOI 5, 4 h) with mCerulean-producing *L. pneumophila* JR32 (pNP99), clicked with DIBO594 dye, fixed, and analyzed by confocal microscopy. Scale bars, 3 µm. Fluorescence intensity profiles were generated for GBP-GFP or P4C-GFP and DIBO594 dye using the RGB profile from ImageJ.

The GBP-GFP fusion construct is biologically active upon ectopic production (**Fig. 4AB**), and thus, we sought to test where the fusion protein localizes in the cell. In uninfected *D. discoideum* GBP-GFP co-localized with calnexin-mCherry, indicating that the large GTPase resides in the ER (**Fig. 5B**). To further assess the localization of GBP, we used dually labeled *D. discoideum* producing GBP-GFP and either the ER marker calnexin-mCherry or the LCV/PtdIns(4)*P* marker P4C-mCherry. Upon infection of the dually labeled *D. discoideum* strains with mCerulean-producing *L. pneumophila* JR32, GBP-GFP co-localized with calnexin-mCherry but not with P4C-mCherry (**Fig. 5CD**, **Fig. S5**). Thus, GBP localizes to the ER at LCV-ER MCS but not to the PtdIns(4)*P*-positive LCV membrane. This localization pattern was observed in *L. pneumophila*-infected *D. discoideum* at 2-12 h p.i., regardless of whether the amoebae were treated with 10 µM LAI-1 or not (**Fig. 5CD**, **Fig. S5**). Taken together, GBP-GFP localizes to the ER but not to the LCV membrane throughout an infection with *L. pneumophila*, and the localization is not affected by LAI-1.

Next, we employed clickable azido-LAI-1 to further characterize the localization of GBP-GFP and to test whether the absence of GBP in the Δ*gnbp* mutant strain affects LAI-1 localization (**Fig. 5E**). In *D. discoideum* Ax2 infected with mCerulean-producing *L. pneumophila* JR32, GBP-GFP co-localized with azido-LAI-1 around LCVs. As expected, GBP-GFP did not localize around intracellular *M. marinum*, which does not reside in an ER-associated compartment (**Fig. S4B**). Moreover, azido-LAI-1 co-localized with the LCV marker P4C-GFP in *D. discoideum* Ax2 (**Fig. 3A**) as well as in Δ*gnbp* mutant amoebae (**Fig. 5E**). Hence, LAI-1 co-localizes with GBP-GFP and with P4C-GFP independently of GBP.

### LAI-1 reduces the size of GBP-positive LCVs

Within the first two hours of *L. pneumophila* infection, LCVs expand in size and are remodeled, likely reflecting the formation of a replication-permissive compartment (Vormittag *et al*., 2023). To assess the role of LAI-1 for LCV size expansion and remodeling, we used dually labeled *D. discoideum* producing GBP-GFP and AmtA-mCherry. Prior to an infection with mCerulean-producing *L. pneumophila* JR32 or Δ*icmT*, the amoebae were treated with 10 µM LAI-1 or not, and the LCV area was quantified by confocal microscopy (**Fig. 6A**, **Fig. S6**). Interestingly, throughout the infection (2-8 h p.i.), treatment with LAI-1 resulted in a significantly smaller size of LCVs harboring wild-type *L. pneumophila* (**Fig. 6B**), but did not affect phagosomes harboring Δ*icmT* mutant bacteria (**Fig. 6C**). Analogously, treatment of *D. discoideum* producing P4C-mCherry with 10 µM LAI-1 or azido-LAI-1 significantly reduced the size of LCVs harboring mCerulean-producing wild-type *L. pneumophila* (**Fig. S1C**). Taken together, treatment with LAI-1 or azido-LAI-1 resulted in significantly smaller LCVs harboring *L. pneumophila* wild-type but not Δ*icmT* mutant bacteria, suggesting that LAI-1 impairs LCV remodeling at early stages of infection.

**Fig. 6.**
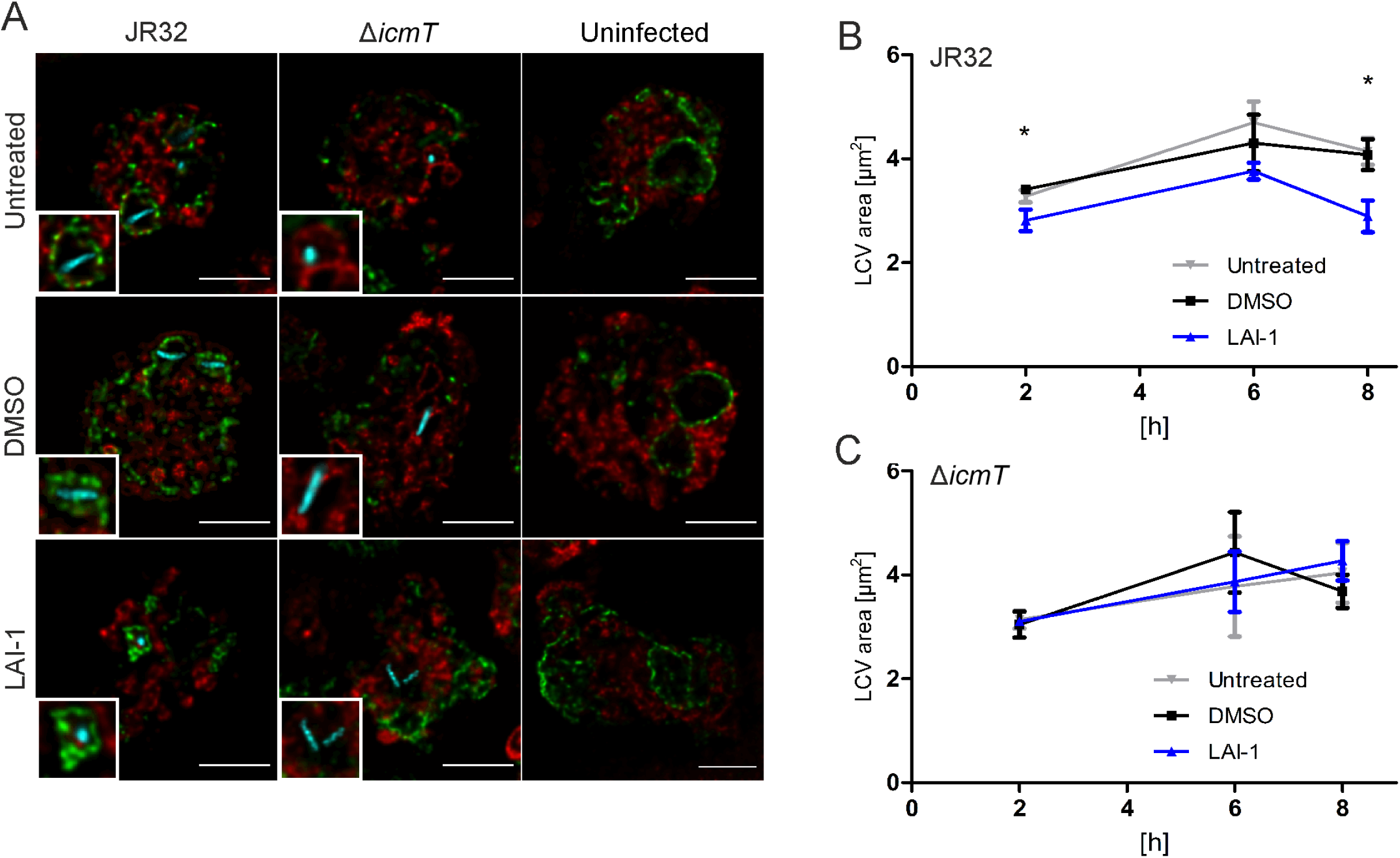
LAI-1 co-localizes with GBP and reduces the size of GBP-positive LCVs. (**A**) Dually labeled *D. discoideum* Ax2 producing GBP-GFP and AmtA-mCherry (pDM1044-AmtA-mCherry) was left untreated or treated with LAI-1 (10 µM, 1 h) or DMSO (solvent control), infected (MOI 5, 8 h) with mCerulean-producing *L. pneumophila* JR32 or Δ*icmT* (pNP99) and analyzed by confocal microscopy. Scale bars, 5 μm. The area of LCVs containing (**B**) strain JR32 or (**C**) strain Δ*icmT* was quantified using ImageJ software. Data shown are means and standard deviations of biological triplicates (Student’s t-test; *, *p* ≤ 0.05).

### LAI-1-dependent LCV size remodeling involves GBP

Since GBP is implicated in LAI-1-dependent inhibition of *D. discoideum* migration (**Fig. 4C**), we sought to assess whether GBP also plays a role in LAI-1-dependent LCV remodeling. To address this question, we used dually labeled *D. discoideum* or Δ*gnbp* mutant amoebae producing calnexin-GFP and P4C-mCherry. Prior to an infection with mCerulean-producing wild-type *L. pneumophila*, the amoebae were treated with 10 µM LAI-1 or not, and the LCV area was quantified by confocal microscopy (**Fig. 7A**, **Fig. S7A**). Throughout the infection (2-20 h p.i.), the treatment of the parental *D. discoideum* strain with LAI-1 resulted in a significantly smaller size of calnexin-GFP-positive LCVs (**Fig. 7B**) and P4C-positive LCVs (**Fig. 7C**). Intriguingly, however, the LCV size reduction by LAI-1 did no longer occur in *D. discoideum* Δ*gnbp*. Accordingly, GBP is implicated in LAI-1-dependent LCV size remodeling, analogously to LAI-1-dependent cell migration inhibition.

**Fig. 7.**
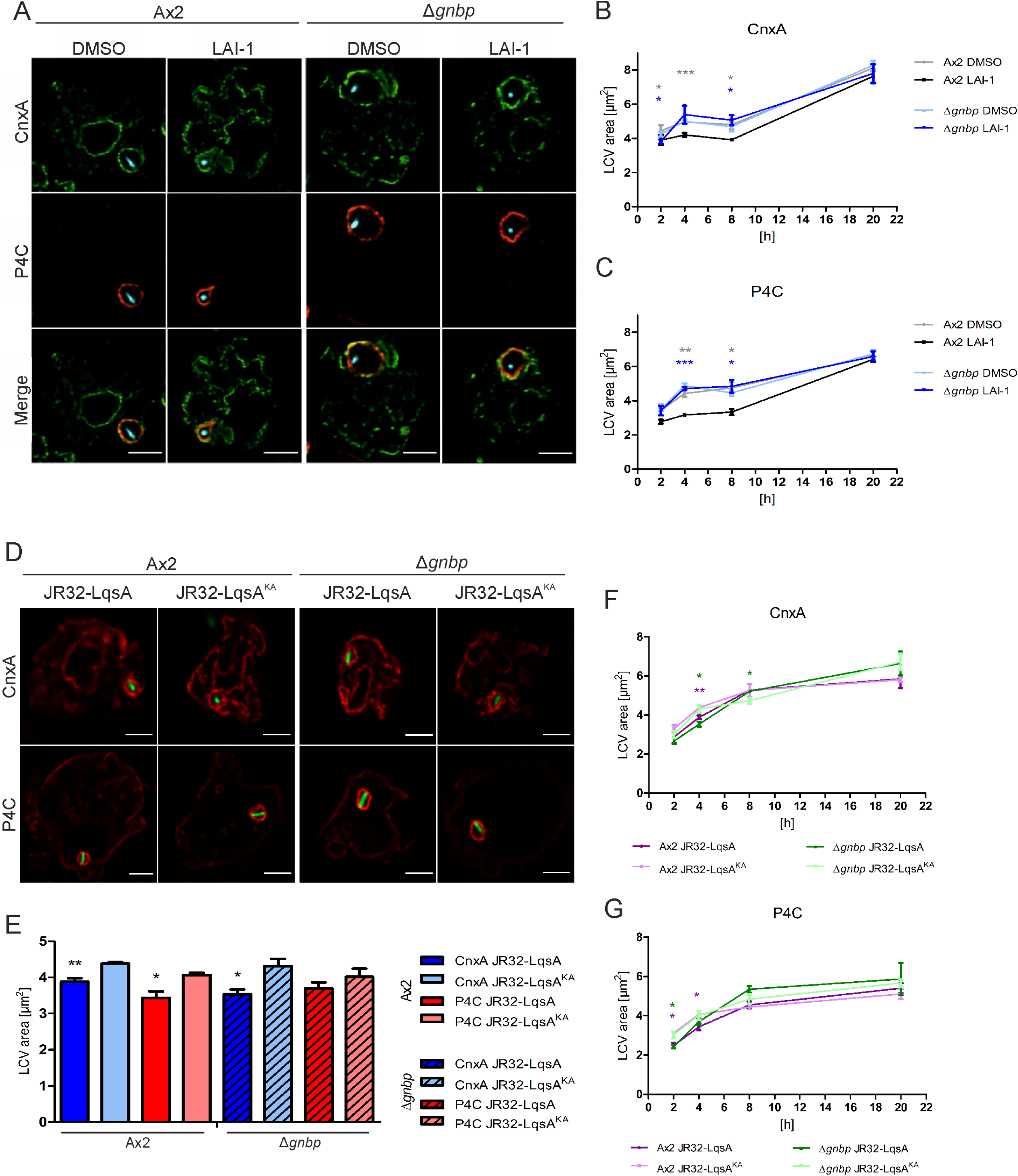
LAI-1-dependent LCV remodeling requires GBP. (**A-C**) Dually labeled *D. discoideum* Ax2 or Δ*gnbp* producing calnexin (CnxA)-GFP (pAW016) and P4C-mCherry (pWS032) was treated with LAI-1 (10 µM, 1 h), or DMSO (solvent control), infected (MOI 5, 4 h) with mCerulean-producing *L. pneumophila* JR32 (pNP99), fixed and analyzed by confocal microscopy. Scale bars, 3 µm. (**D-G**) *D. discoideum* Ax2 or Δ*gnbp* producing CnxA-mCherry (pAW012) or P4C-mCherry (pWS032) was infected (MOI 5) for (**D**, **E**) 4 h or (**F**, **G**) 2-20 h with GFP-producing *L. pneumophila* JR32 harboring pMF16 (P*_6SRNA_*-*lqsA*) or pMF17 (P*_6SRNA_*-*lqsA*^K258A^), fixed and analyzed by confocal microscopy. Scale bars, 3 µm. The area of LCVs positive for (**E**, **F**) CnxA or (**E, G**) P4C was quantified using ImageJ software. Data shown are means and standard deviations of biological triplicates (Student’s t-test; *, *p* ≤ 0.05; **, *p* ≤ 0.01; ***, *p* ≤ 0.001).

We also tested whether the effects of synthetic LAI-1 on LCV size remodeling are observed with endogenously produced LAI-1. To this end, we used GFP-tagged *L. pneumophila* JR32 overproducing wild-type LqsA or, as a negative control, the catalytically inactive mutant LqsA^K258A^ under control of the strong P*_6SRNA_* promoter (Fan *et al*., 2023). The *D. discoideum* parental strain Ax2 or Δ*gnbp* mutant amoebae producing calnexin-mCherry or P4C-mCherry were infected with these bacterial strains, and the LCV area was quantified by confocal microscopy (**Fig. 7D**, **Fig. S7B**). In the parental *D. discoideum* Ax2 strain, calnexin-positive (**Fig. 7EF**) LCVs as well as PtdIns(4)*P*-positive (**Fig. 7EG**) LCVs harboring *L. pneumophila* overproducing LqsA were significantly smaller than LCVs harboring *L. pneumophila* overproducing LqsA^K258A^. These findings indicate that endogenously produced LAI-1 affects LCV remodeling. In Δ*gnbp* mutant amoebae, LqsA-dependent size remodeling no longer occurred with PtdIns(4)*P*-positive LCVs (**Fig. 7E**) but still happened with calnexin-positive LCVs (**Fig. 7E**). In summary, LAI-1- and LqsA affect LCV size remodeling, and a role for GBP in this process is more pronouncedly observed with synthetic LAI-1 than with endogenously produced LAI-1.

In mammalian cells, GBPs compromise the integrity of pathogen vacuoles, thereby contributing to the antimicrobial effect of this protein family (Kutsch & Coers, 2021; Tretina *et al*., 2019). To test whether LAI-1 and/or DdGBP affect LCV integrity, we used *D. discoideum* producing cytosolic mCherry, which is excluded from intact vacuoles but enters compromised pathogen vacuoles (Koliwer-Brandl *et al*, 2019). Dually labeled *D. discoideum* or Δ*gnbp* producing cytosolic mCherry and P4C-GFP were treated with 10 µM LAI-1 or not, infected with mCerulean-producing *L. pneumophila* JR32 and analyzed by confocal microscopy (**Fig. S8**). This approach revealed that LCV integrity is not compromised in either the parental *D. discoideum* Ax2 strain or the Δ*gnbp* mutant at 4 h p.i. Taken together, neither LAI-1 nor GBP affect LCV integrity, suggesting specific roles for LAI-1 and GBP in LCV remodeling and intracellular growth of *L. pneumophila* rather than the mere disruption of the LCV architecture.

## Discussion

In this study, we investigated the role of the single member of the GBP family of large GTPases in *D. discoideum* for LAI-1-dependent inter-kingdom signaling and intracellular replication of *L. pneumophila*. We show that LAI-1 and clickable derivatives impair *D. discoideum* migration, modulate LCV size and localize to ER-LCV MCS (**Fig. 2, 3, 6**), and that GBP impedes *D. discoideum* migration (**Fig. 4**), restricts intracellular growth of *L. pneumophila* (**Fig. 5**) and localizes to ER-LCV MCS (**Fig. 5**). Intriguingly, LAI-1-dependent inhibition of *D. discoideum* migration (**Fig. 4**) and LCV expansion/remodeling involves GBP (**Fig. 7**) without compromising the integrity of LCVs (**Fig. S8**). Current insights on LAI-1- and GBP-dependent inter-kingdom signaling between *L. pneumophila* and host cells are summarized in a working model (**Fig. 1A**).

Intriguingly, DdGBP was previously identified in the proteome of LCVs purified from *L. pneumophila*-infected amoebae (Hoffmann *et al*., 2014a). We validated and corroborated this finding by demonstrating that GBP-GFP is functional (**Fig. 4AB**) and indeed localizes to the ER in uninfected *D. discoideum* and to LCV-ER MCS in *L. pneumophila*-infected amoebae (**Fig. 5B-D**). The LCV-ER MCS form within 1-2 h after the uptake of *L. pneumophila* and are maintained throughout the infection (Vormittag *et al*., 2023b).

Previously, LAI-1 has been shown to inhibit the migration of *D. discoideum*, mouse macrophages and human epithelial cells (Simon *et al*., 2015). In epithelial cells, LAI-1-dependent cell migration inhibition requires the scaffold protein IQGAP1, the small GTPase Cdc42 and the Cdc42-specific guanine nucleotide exchange factor ARHGEF9, but not other modulators of Cdc42, or the small GTPases RhoA, Rac1 or Ran (Simon *et al*., 2015). In the eukaryotic signal transduction pathway triggered by LAI, GBP likely participates upstream of IQGAP1, Cdc42 and ARHGEF9. It remains to be elucidated, which of the 7 human GBP paralogue(s) is implicated in LAI-1-dependent migration inhibition.

*D. discoideum* GBP restricts the intracellular replication of *L. pneumophila* (**Fig. 5A**), without compromising LCV integrity (**Fig. S8**). LCV rupture takes place only late in the *L. pneumophila* infection cycle (> 48 h p.i.), followed by only a very short stay in the host cytoplasm before lysis of the amoeba within minutes (Striednig *et al*., 2021). Given the very late and short exposure of *L. pneumophila* to the host cytoplasm, it is unlikely that DdGBP restricts bacterial growth by directly targeting and killing the pathogen. In agreement with this notion, we did not observe GBP-GFP localizing to cytosolic *L. pneumophila*. Rather, GBP might affect the formation and/or maturation of the LCV. Hence, it appears that by affecting the expansion and remodeling of LCVs, GBP impairs the formation of a replication-permissive compartment and thus might restrict intracellular growth of *L. pneumophila*. In contrast, GBP does not affect intracellular growth of *M. marinum* (**Fig. S4**). This difference might be explicable by the fact that the LCV associates with ER, while the *Mycobacterium*-containing vacuole (MCV) does not. Hence, the mechanism of intracellular growth restriction by GBP might specifically involve the ER.

The complex process of LCV formation and maturation occurs within 1-2 h after uptake of *L. pneumophila* and involves a PI lipid conversion from PtdIns(3)*P* to PtdIns(4)*P* (Weber *et al*., 2018), ER acquisition (Lu & Clarke, 2005; Ragaz *et al*, 2008) and pathogen vacuole size expansion/remodeling (Vormittag *et al*., 2023). Synthetic LAI-1 localizes to the ER at LCV-ER MCS as well as to the PtdIns(4)*P*-positive LCV membrane (**Fig. 2, 3**). Intriguingly, LAI-1 specifically localizes to LCVs harboring wild-type *L. pneumophila* but not to the membrane of AmtA-positive vacuoles harboring avirulent *L. pneumophila* Δ*icmT* (**Fig. 6**). Therefore, specific LCV membrane components might determine LAI-1 acquisition and accumulation. Such factors are currently unknown but might comprise lipids and/or proteins.

Synthetic LAI-1 inhibits the expansion/remodeling of LCVs (**Fig. 6**) in a GBP-dependent manner during the initial 8 h of infection (**Fig. 7A-C**). Perhaps due to effects of LAI-1 on early LCV formation events, synthetic LAI-1 might be more effective than (continuously) released endogenous LAI-1 synthesized by overproduced autoinducer synthase LqsA (**Fig. 7D-G**). Alternatively, the relatively high concentration of 10 µM synthetic LAI-1 might not be reached upon production of the autoinducer by *L. pneumophila*, and/or the solubility and bioavailability of synthetic LAI-1 and endogenous LAI-1 might differ. Notably, endogenous LAI-1 is associated with and released through OMVs by *L. pneumophila* (Fan *et al*., 2023).

The LAI-1-dependent size reduction of LCVs is relatively small – untreated LCVs were ca. 4-5 µm^2^, compared to LAI-1-treated LCVs which were ca. 3-4 µm^2^ at 8 h p.i. – and thus likely reflects a structural remodeling of the pathogen vacuole rather than a substantial LCV expansion. A massive expansion of the pathogen vacuole occurs at later infection time points (> 8 h p.i.), likely through the interception of (anterograde and retrograde) vesicular trafficking between the ER and the Golgi apparatus (Kagan & Roy, 2002; Nagai *et al*, 2002; Robinson & Roy, 2006), and through the fusion of Golgi-derived vesicles (Weber *et al*., 2018).

Mammalian GBPs function as PRRs (Kirkby *et al*., 2023), which bind conserved bacterial structures such as LPS of Gram-negative bacteria, and thus, contribute to pathogen detection and elimination (Kutsch *et al*., 2020; Santos *et al*., 2020). Analogously, *D. discoideum* GBP might recognize conserved bacterial patterns such as LPS or certain classes of bacterial low molecular weight molecules. Accordingly, DdGBP might represent an evolutionarily ancient PRRs, which has evolved to bind and detect small lipophilic molecules such as the lipid A anchor of LPS or hydrophobic signaling molecules such as the aliphatic α-hydroxyketones LAI-1 (3-hydroxypentadecane-4-one) and *V. cholerae* CAI-1 (3-hydroxytridecane-4-one). It is currently unknown whether DdGBP directly binds LAI-1 and what effects this binding might have on a molecular and cellular level.

Overall, our findings agree with the notion that similarly to mammalian cells, the *D. discoideum* GBP is implicated in the recognition of and/or defense against intracellular pathogens, and therefore, functions as an antimicrobial compound. However, since the protozoan amoebae do not produce cytokines, caspases, or their activation platforms like the inflammasome, the output of GBP-dependent pathogen detection by amoebae does not involve pyroptotic or apoptotic cell death, and thus, is clearly different from mammalian cells. Given that DdGBP restricts the intracellular replication of *L. pneumophila* but not *M. marinum* (**Fig. S4A**) and GBP-GFP localizes to LCVs but not MCVs (**Fig. S4B**), the ER-association of GBP might underly the specificity of bacterial killing.

In summary, LAI-1-dependent inter-kingdom signaling of *L. pneumophila* comprises cell migration inhibition and cytoskeleton remodeling, as well as LCV expansion and dynamics. LAI-1 is detected and taken up by eukaryotic host cells by unknown mechanisms. The single *D. discoideum* GBP family large GTPase restricts intracellular growth of *L. pneumophila*, localizes to LCV-ER contact sites and is implicated in LAI-1-dependent LCV remodeling. Thus, LAI-1 links small molecule inter-kingdom signaling and GBP-dependent cell autonomous immunity, as outlined in the working model (**Fig. 1A**). These results collectively suggest a novel mechanism of inter-kingdom signaling mediated by LAI-1 and GBP, shedding light on the intricate pathogen-host interactions between *L. pneumophila* and host cells. Further studies will elucidate the pathways underlying the inter-kingdom detection of and response to LAI-1 by eukaryotic host cells.

## Materials and Methods

### Bacteria, *D. discoideum*, and Δ*gnbp* mutant strain

The bacterial strains and cell lines used in this study are listed in **Table 1**. *L. pneumophila* was grown for 3 days on charcoal yeast extract (CYE) agar plates (Feeley *et al*, 1979), with or without chloramphenicol (Cam; 10 µg/ml) at 37°C. Bacterial colonies were used to inoculate liquid cultures on a wheel (starting concentration OD_600_ of 0.1, 80 rpm) in *N*-(2-acetamido)-2-aminoethanesulfonic acid (ACES)-buffered yeast extract (AYE) medium (Horwitz, 1983) and grown for approximately 21 h at 37°C to an early stationary phase (2 × 10^9^ bacteria/ml), with Cam (5 µg/ml) added to maintain plasmids if required.

*V. cholerae* strain MM920 was cultured overnight at 30°C in LB broth supplemented with tetracycline (Tet; 5 μg/ml) prior to an experiment. *V. cholerae* MM920 lacks the sensor kinase gene *luxQ* and the autoinducer synthase gene *cqsA*, and therefore, does not respond to AI-2, and does not produce but responds to the α-hydroxyketone compounds CAI-1 and LAI-1. Strain MM920 harbors plasmid pBB1, which contains the *luxCDABE* luciferase operon of *V. harveyi* and produces light upon detection of CAI-1 and LAI-1.

*D. discoideum* strains were grown at 23°C in HL5 medium without glucose (ForMedium) supplemented with D-maltose (Roth). The amoebae were grown in T75 flasks and split every second day. *D. discoideum* was transformed and selected with geneticin (G418; 20 μg/ml), hygromycin (Hyg; 50 μg/ml), or blasticidin S (Bls; 5 µg/ml) as described previously (Knobloch *et al*, 2020; Koliwer-Brandl *et al*., 2019; Weber & Hilbi, 2014; Weber *et al*., 2014).

**Table 1.**
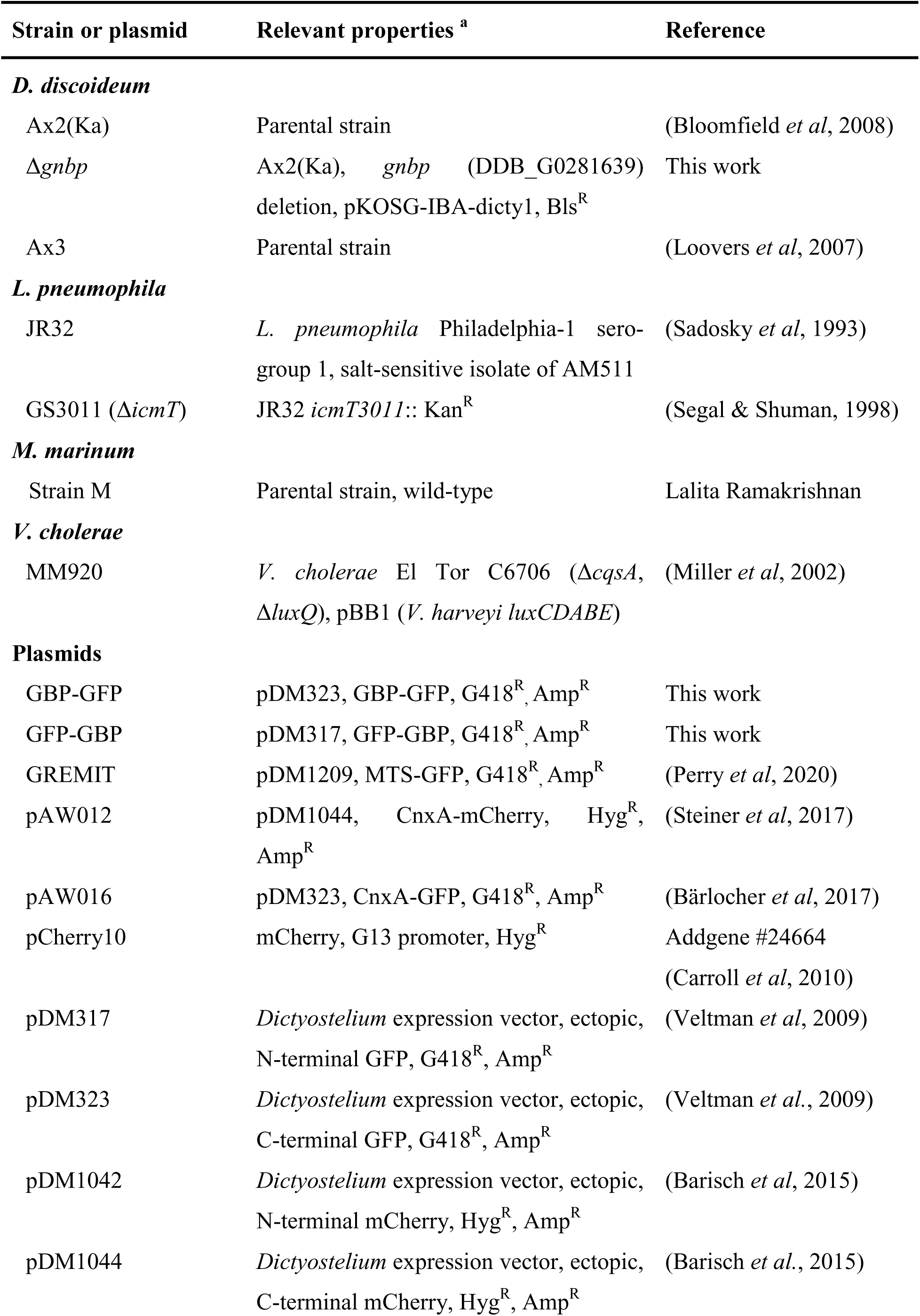
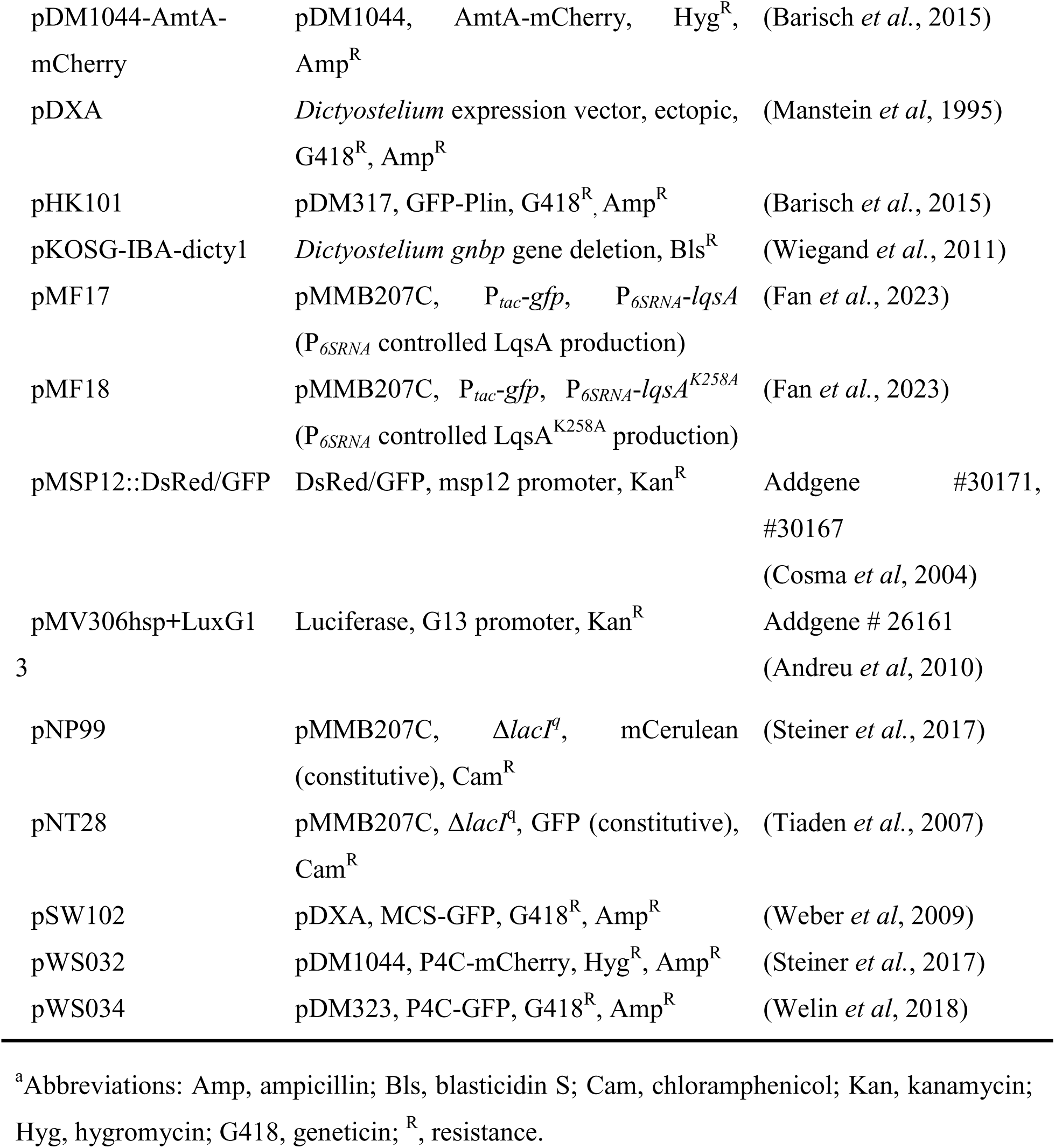
Cells, bacterial strains, and plasmids used in this study.

The *D. discoideum gnbp* gene (DDB_G0281639) was deleted in *D. discoideum* Ax2(Ka) by double homologous recombination using plasmid pKOSG-IBA-dicty1, yielding strain Δ*gnbp* (**Table 1**). To this end, strain Ax2(Ka) was grown in axenic conditions at 22°C in HL5c medium (Formedium) supplemented with 100 U/ml penicillin and 100 µg/ml streptomycin (Invitrogen). The Δ*gnbp* deletion mutant strain was generated by homologous recombination following a one-step cloning in pKOSG-IBA-dicty1 as previously described (Wiegand *et al*, 2011). The GFP-GBP or GBP-GFP constructs were generated by cDNA library PCR-amplification of the *gnbp* open reading frame using the oligonucleotides indicated (**Supplementary Table S1**), flanked by BglII / XbaI restriction sites and inserted into the pDM317 or pDM323 backbones digested with BglII / SpeI (XbaI complementary). The GFP-GBP construct was used throughout this study, while the GBP-GFP construct appeared to produce unspecific GFP aggregates in *D. discoideum*.

### *V. cholerae* LAI-1 reporter assay

The *V. cholerae* strain MM920 was inoculated in LB liquid medium supplemented with Tet (5 μg/ml) and incubated for 18 h at 37 °C. The overnight culture was diluted to an OD_600_ of 0.25 with fresh medium supplemented with synthetic LAI-1 or derivatives (1-50 µM), or DMSO as solvent control. The mixtures were then transferred to a 96-well plate (Chemie Brunschwig AG), and luminescence intensity (bottom read) was measured every 0.5 h for 8 to 10 h at 30°C using a Biotek Cytation 5 microplate reader with continuous orbital shaking. Images were captured after 4 to 5 h incubation (when bioluminescence intensity usually reached maximum levels) using the FluorChem SP imaging system (Alpha-InnoTec) with an exposure time of 15 min.

### Synthesis of LAI-1, azido-LAI-1, and diazirine-axido-LAI-1

(*S*)-LAI-1 was synthesized as described (Simon *et al*., 2015) and is referred to as “LAI-1” throughout the manuscript. Azido-(*S*)-LAI-1 and diazirine-azido-(*S*)-LAI-1 are referred to as “azido-LAI-1” and “diazirine-LAI-1”, respectively, and were synthesized as follows (**Fig. S9, Fig. S10**): The chemical synthesis of clickable azido-LAI-1 (**9**) and bifunctional photoreactive diazirine-LAI-1 (**22**) started with commercially available (*S*)-2-hydroxybutyric acid **1**. In contrast to the previously described synthesis of (*S*)-LAI-1 (Simon *et al*., 2015), the carboxylic acid was first converted to the corresponding Weinreb amide **2** (Fink *et al*., 2021), and then the alcohol was protected as a silyl ether **3**. This modification was needed as the original synthesis route yielded only poor results during scaling up. For the synthesis of azido-LAI-1, the Weinreb amide **3** was reacted with the acetal-protected bromide **4** via Grignard reaction to form the ketone **5**. The hydroxyl group was released via acid-catalyzed cleavage in a protic medium and subsequently converted into bromide **7** using Appel reaction conditions. The bromide was converted to the azide (Lang *et al*, 2020), and deprotection of the silyl ether with TBAF yielded azido-LAI-1 (**9**).

The synthesis of diazirine-LAI-1 was also initiated via a Grignard reaction with the Weinreb amide **3**, but an acetal-protected bromide **10** with a shorter chain length was used to allow subsequent functionalization of the side chain. The terminal acetal **11** was acid-catalyzed and cleaved to obtain the free alcohol **12**. To facilitate the upcoming Grignard reaction and diazirine synthesis, the carbonyl group had to be protected as a 1,3-dioxolane **13** to ensure selective conversion. Camphorsulfonic acid was required for this step, as stronger acids such as *para*-toluenesulfonic acid (Yang *et al*, 2000) led to complete decomposition of the reactant, while weaker acids such as pyridinium *para*-toluenesulfonate resulted in no conversion at all. The primary alcohol **13** was oxidized to carboxylic acid **14** by pyridinium dichromate (Corey & Schmidt, 1979), which was directly converted into the Weinreb amide **15**. The ketone **17**, containing a carbonyl group in the middle of the side chain, was formed through a Grignard reaction of the Weinreb amide **15** with the acetal-protected bromide **16**. The carbonyl group was transformed in methanolic ammonia solution with hydroxylamine-*O*-sulfonic acid (HOSA) into the relatively unstable diaziridine, which was then directly oxidized with iodine to form diazirine **18** (Procacci *et al*, 2018). The diaziridine synthesis was also attempted in liquid ammonia (Wang *et al*, 2017), but it was unsuccessful due to insolubility. Subsequently, the carbonyl group was very gently deprotected via a Lewis acid-catalyzed transacetalization with acetone and a catalytic amount of iodine to avoid cleaving the silyl ether (Sun *et al*, 2004). The acetal deprotection to the primary alcohol **19** and the following reaction steps to diazirine-LAI-1 (**22**) were carried out analogously to the synthesis of azido-LAI-1 (**9**).

### Click chemistry of azido-LAI-1

*D. discoideum* strains producing GFP-labeled organelle markers were seeded in a 12-well plate (5 × 10^5^ cells/well) in HL5 medium and cultured overnight at 23°C. On the day of infection, the amoebae were treated with 10 µM azido-LAI-1 or DMSO (solvent control) for 1 h. Subsequently, the amoebae were infected (MOI 5) with mCerulean-producing *L. pneumophila* JR32 (pNP99), centrifuged (450 × *g*, 10 min; room temperature, RT) and incubated at 25°C for 1 h. The infected amoebae were washed three times with HL5 medium and further incubated for the indicated times. 30 min before stopping the infection at the specific time point, 10 µM of clickable DIBO594 dye (ThermoScientific, C10407) was added and further incubated at 25°C. Cells were collected from the 12-well plates and centrifuged (500 × *g*, 5 min, RT). The supernatant was discarded, the cell pellet resuspended in HL5 medium and centrifuged again (500 × *g*, 5 min, RT). The samples were fixed with 4% PFA (Electron Microscopy Sciences; 30 min, RT), washed twice with PBS, transferred to an 18-well μ-slide dish (Ibidi) and immobilized by adding 0.5% agarose in PBS.

For imaging of the samples, a confocal laser scanning microscope Leica TCS SP8 X CLSM (HC PL APO CS2, objective 63×/1.4-0.60 oil; Leica Microsystems) was used, with a scanning speed of 100 Hz and bi-directional laser scan. Acquisition was performed with a pixel/voxel size close to the instrument’s Nyquist criterion of 43 × 43 × 130 nm (xyz). Deconvolution of the images was performed with Huygens professional version 19.10 software (Scientific Volume Imaging, http://svi.nl) using the CMLE algorithm, set to 10-20 iterations and 0.05 quality thresholds. The colocalization of clickable LAI-1 with different cellular markers was quantified by using ImageJ plugin “Coloc 2” obtaining Pearson’s correlation coefficients.

### Photo-crosslinking of diazirine-LAI-1

One day before the experiment, *D. discoideum* Ax2 was seeded (5 × 10^5^ cells per well) in 12-well plates and cultured overnight at 23°C. 1 h before infection, the amoebae were treated with 10 µM diazirine-LAI-1 or azido-LAI-1. The amoebae were infected (MOI 5) with mCerulean-producing *L. pneumophila* JR32 (pNP99), centrifuged (450 × *g*, 10 min, RT) and incubated at 25°C for 1 h. Subsequently, infected cells were washed three times with HL5 medium to remove extracellular bacteria, the medium was supplemented again with the respective LAI-1 derivative and further incubated at 25°C for the time indicated. At 8 h p.i., given samples were UV-irradiated for 5 min on ice using a 40 W Hg lamp (8 W, 5 bulbs), operated at 1000 W at a distance of ca. 35 cm from the light source. After 24 h p.i., cells (including supernatant) were collected by vigorous pipetting, centrifuged (500 × *g*, 5 min, RT) and fixed with 4% PFA (30 min, RT). After fixation, the amoebae were washed twice with PBS, transferred to an 18-well μ-slide dish (Ibidi) and immobilized by adding 0.5% agarose in PBS to the wells. For imaging of the samples, a confocal laser scanning microscope Leica TCS SP8 X CLSM was used as described above.

### Single amoeba tracking

*D. discoideum* strains Ax3, Ax2, Δ*gnbp*, Δ*gnbp*/GFP or Δ*gnbp*/GBP-GFP were seeded at a density of 2×10^4^ cells/well into an 8-well μ-slide dish (Ibidi) and incubated for 3-4 h in HL5 medium to allow attachment. The medium was then replaced by MB medium, and the amoebae were incubated at 23°C for 1 h before imaging. During microscopy, three fields of interest were randomly selected for each sample and recorded continuously for 2 h with 2 min time interval. Image analyses were performed using ImageJ and the Chemotaxis and Migration Tool version 2.0 (Ibidi).

### Chemotaxis migration assay

Under-agarose migration assays with *D. discoideum* were performed as described (Laevsky & Knecht, 2001; Simon *et al*, 2014). The day before the assay, GFP-producing *D. discoideum* Ax2 or Δ*gnbp* (pDM317) were seeded into 6-well plates in HL5 medium (1 × 10^6^ cells/well), and microscopy dishes (μ-Dish, 35 mm, Ibidi) were filled with a mixture of melted agarose in SM medium [10 g bacteriological peptone (Oxoid), 1 g Bacto yeast extract (BD Biosciences), 1.9 g KH_2_PO_4_, 0.6 g K_2_HPO_4_, 0.43 g MgSO_4_, 10 g glucose per liter, pH 6.5]. After solidification, 3 parallel slots of 2 × 4 mm (for cells and chemo-attractant solution) were manually cut 5 mm apart into the agarose. On the day of the experiment, the amoebae were washed once with MB medium [14 g bacteriological peptone (Oxoid), 7 g Bacto yeast extract (BD Biosciences), 4.26 g MES (Sigma-Aldrich) per liter, pH 6.9] and kept for 1 h in MB medium. During this period, LAI-1 or DMSO were added at the concentrations indicated. In parallel, the dishes were prepared by adding the chemo-attractant solution, 1 mM folic acid (Sigma-Aldrich) in SM medium, into the central slot 30 min before the cell suspensions were filled into the neighboring slots. After 2 washing steps with MB medium (450 × *g*, 10 min), the amoebae were detached by scratching into 500 μl MB, and 30 μl of the cell suspension was filled into the slots. The dishes were incubated for 4 h at 23°C to let the amoebae migrate. The migration was tracked using a Leica TCS SP8 X CLSM microscope as described above.

### Uptake and intracellular replication of *L. pneumophila*

Uptake of GFP-producing *L. pneumophila* JR32 or Δ*icmT* by *D. discoideum* Ax2 or Δ*gnbp* was assessed by flow cytometry. To this end, exponentially growing *D. discoideum* was seeded onto a 24-well plate (1 × 10^5^ cells/well) in HL5 medium and cultured overnight at 25°C. On the day of the infection, the amoebae were treated with 10 µM LAI-1 or DMSO (solvent control) for 1 h and infected (MOI 50) with the *L. pneumophila* strains harboring plasmid pNT28. *L. pneumophila* strains were grown for 21 h in AYE/Cam, diluted in HL5 medium, centrifuged onto the cells (450 × *g*, 10 min, RT) and further incubated for 30 min at 25°C, followed by washing three times with HL5 medium to remove extracellular bacteria. Infected *D. discoideum* were detached by vigorously pipetting, and 2 × 10^4^ amoebae per sample were analyzed using a LSR II Fortessa cell analyzer (Becton Dickinson, Palo Alto, United States). Scatter plot gating was based on uninfected amoebae, and GFP fluorescence intensity was quantified using FlowJo software.

Intracellular growth of GFP-producing *L. pneumophila* JR32 or Δ*icmT* in *D. discoideum* Ax2 or Δ*gnbp* was assessed by fluorescence increase (relative fluorescence units, RFU). To this end, *D. discoideum* amoebae were seeded (2 × 10^4^ cells per well) in 96-well culture-treated plates (ThermoFisher) and cultured in HL5 medium overnight at 23°C. Amoebae were treated with 10 µM LAI-1 or DMSO for 1 h and infected (MOI 1) with the *L. pneumophila* strains harboring plasmid pNT28. *L. pneumophila* strains were grown for 21 h in AYE medium, diluted in MB medium, centrifuged onto the cells (450 × g, 10 min, RT) and incubated for 1 h at 25°C. Subsequently, the medium was exchanged with fresh MB medium (supplemented with LAI-1 or DMSO) and further incubated for the time indicated at 25°C. GFP fluorescence was measured every two days using a BioTek Cytation 5 microplate reader (Agilent Technologies).

### Quantification of LCV area sizes in *D. discoideum*

Dually fluorescence-labeled *D. discoideum* strains were grown in exponential phase in HL5 medium containing geneticin (G418, 20 μg/ml) and/or Hyg (50 μg/ml). One day before the experiment, the amoebae were seeded (5 × 10^5^ cells per well) in 12-well plates and cultured overnight at 23°C. 1 h before infection, the amoebae were treated with 10 µM LAI-1, LAI-1 derivative or DMSO (solvent control). The amoebae were infected (MOI 5) with mCerulean-producing *L. pneumophila* JR32 (pNP99), centrifuged (450 × *g*, 10 min, RT) and incubated at 25°C for 1 h. Subsequently, infected cells were washed three times with HL5 medium to remove extracellular bacteria, the medium was supplemented again with the respective LAI-1 derivative or DMSO and further incubated at 25°C for the time indicated. At given infection time points, cells (including supernatant) were collected by vigorous pipetting, centrifuged (500 × *g*, 5 min, RT) and fixed with 4% PFA (30 min, RT). After fixation, the amoebae were washed twice with PBS, transferred to an 18-well μ-slide dish (Ibidi) and immobilized by adding 0.5% agarose in PBS to the wells. For imaging of the samples, a confocal microscope Leica TCS SP8 X CLSM was used as described above.

### Statistical methods

Each experiment was independently performed at least three times and representative images are shown. All statistical analyses were performed using GraphPad Prism (www.graphpad.com). The two-tailed Student’s *t*-test (Mann-Whitney test, no assumption of Gaussian distributions) was used to show significant differences between samples and control. Significances are indicated in the figures as follows: *, **, *** or **** to indicate probability values of less than 0.05, 0.01, 0.001 and 0.0001, respectively. The value of “n” represents the number of biological replicates performed or the number of analyzed cells/LCVs per condition.

## Abbreviations

Icm/Dot: intracellular multiplication/defective organelle trafficking
c-di-GMP: cyclic di-guanosine monophosphate
DBCO: dibenzocyclooctyne
h/mGBP: human/mouse guanylate binding protein
IFN-γ: interferon-γ
LAI-1: *Legionella* autoinducer-1
LCV: *Legionella*-containing vacuole
Lqs: *Legionella* quorum sensing
LvbR: *Legionella* virulence and biofilm regulator
GBP: guanylate binding protein
GFP: green fluorescent protein
OMVs: outer membrane vesicles
T4SS: type IV secretion system.

## Data availability

All data are contained within the manuscript.

## Acknowledgements

This work was supported by Swiss National Science Foundation (SNF) project grants to HH (31003A_175557, 310030_200706) and TS (310030_169386, 310030_188813) as well as a SNF Sinergia grant to TS (CRSII5_189921). Work in the group of JS was supported by the Deutsche Forschungsgemeinschaft (DFG) within the research training group RTG2581. The authors have no conflict of interest to declare.

## Figure Legends

**Fig. S1. LAI-1 and clickable derivatives promote luminescence of a *Vibrio* reporter strain and LCV size modulation. (A)** The *Vibrio cholerae* α-hydroxyketone reporter strain MM920 was left untreated or treated with the indicated concentrations of synthetic (*S*)-LAI-1, azido-LAI-1, diazirine-LAI-1, or DMSO (solvent control), and luminescence intensity was measured by a plate reader (30°C, 10 h). RLU, relative light units. Data shown are biological triplicates of means and standard deviations. (**B**) *D. discoideum* Ax2 was treated with LAI-1 or clickable azido-LAI-1 (10 µM, 1 h), infected (MOI 5, 4 h) with mCerulean-producing *L*. *pneumophila* JR32 (pNP99), clicked with DIBO594 dye, and analyzed by confocal laser scanning microscopy. Scale bars: 20 μm. (**C**) *D. discoideum* Ax2 producing P4C-mCherry (pWS032) was treated (10 µM, 1 h) with LAI-1, azido-LAI-1, or DMSO (solvent control), infected (MOI 5, 4 h) with mCerulean-producing *L. pneumophila* JR32 (pNP99), fixed and analyzed by confocal microscopy. LCV areas were quantified using ImageJ software. Data shown are means and standard deviations of biological triplicates (Student’s t-test; *, *p* ≤ 0.05; **, *p* ≤ 0.01).

**Fig. S2. LAI-1-dependent migration inhibition of *D. discoideum* involves GBP.** *D. discoideum* Ax2 or Δ*gnbp* producing GFP (pDM317) was left untreated or treated with LAI-1 (1 µM, 5 µM or 10 µM; 1 h) or DMSO (solvent control), and cell migration towards 1 mM folate (4 h) was assessed by under-agarose assay. The white lines represent the edge of the sample wells.

**Fig. S3. *D. discoideum* Δ*gnbp* does not permit replication of *L. pneumophila* Δ*icmT* and does not affect *L. pneumophila* uptake.** (**A**) *D. discoideum* Ax2 or Δ*gnbp* was infected (MOI 1, 10 d) with GFP-producing *L. pneumophila* JR32 or Δ*icmT* (pNT28), and intracellular replication was assessed by RFU. Data shown are means and standard deviations of biological triplicates (Student’s t-test; *, *p* ≤ 0.05; **, *p* ≤ 0.01). (**B**, **C**) *D. discoideum* Ax2 or Δ*gnbp* was treated with LAI-1 (10 µM, 1 h) or DMSO (solvent control), infected (MOI 50, 30 min) with GFP-producing *L. pneumophila* JR32 (pNT28) and analyzed by flow cytometry. Untreated, uninfected amoebae were used for gating. Data shown are (B) counts vs. GFP fluorescence intensity, and (C) percentage of GFP-positive amoebae (means and standard deviations of biological triplicates).

**Fig. S4. GBP does not affect growth of intracellular *M. marinum*.** (**A**) *D. discoideum* Ax2 or Δ*gnbp* was infected (MOI 10) for the time indicated with luciferase-producing *M*. *marinum*, and intracellular growth was assessed by bioluminescence. Means and SEM of biological triplicates are shown for three independent Δ*gnbp* clones (c3, c17, c19); r.l.u., relative light units. (**B**) *D. discoideum* Ax2 producing GBP-GFP was infected (MOI 10, 1.5 h or 24 h) with mCherry-producing *M. marinum*, fixed and analyzed by confocal microscopy. Representative maximum projections of live time-lapse spinning disk confocal images are shown. Scale bars, 10 µm. Images are representative of at least 3 independent experiments.

**Fig. S5. GBP localizes to ER at LCV-ER contact sites.** Dually labeled *D. discoideum* Ax2 producing GBP-GFP and (**A**) calnexin (CnxA)-mCherry (pAW012) or (**B**) P4C-mCherry (pWS032) was left untreated or treated with LAI-1 (10 µM, 1 h), or DMSO (solvent control), infected (MOI 5, 4 h) with mCerulean-producing *L. pneumophila* JR32 (pNP99), fixed, and analyzed by confocal microscopy. Scale bars, 3 µm. Single channels and merge are shown (related to **Fig. 5**).

**Fig. S6. LAI-1 prevents expansion of GBP-positive LCVs.** Dually labeled *D. discoideum* Ax2 producing GBP-GFP and AmtA-mCherry (pDM1044-AmtA-mCherry) was left untreated or treated with LAI-1 (10 µM, 1 h), or DMSO (solvent control), infected (MOI 5, 8 h) with mCerulean-producing *L. pneumophila* JR32 or Δ*icmT* (pNP99) and analyzed by confocal microscopy. Scale bars, 3 μm. Single channels and merge are shown (related to **Fig. 6**).

**Fig. S7. LAI-1-dependent LCV remodeling involves GBP.** (**A**) Dually labeled *D. discoideum* Ax2 or Δ*gnbp* producing calnexin (CnxA)-GFP (pAW016) and P4C-mCherry (pWS032) was treated with LAI-1 (10 µM, 1 h), or DMSO (solvent control), infected (MOI 5, 4 h) with mCerulean-producing *L. pneumophila* JR32 (pNP99), fixed and analyzed by confocal laser microscopy. Scale bars, 3 µm. (**B**) *D. discoideum* Ax2 or Δ*gnbp* producing CnxA-mCherry (pAW012) or P4C-mCherry (pWS032) was infected (MOI 5, 4 h) with GFP-producing *L. pneumophila* JR32 harboring pMF16 (P*_6SRNA_*-*lqsA*) or pMF17 (P*_6SRNA_*-*lqsA*^K258A^), fixed and analyzed by confocal microscopy. Scale bars, 3 µm. Single channels and merge are shown (related to **Fig. 7**).

**Fig. S8. LAI-1 and GBP do not affect LCV integrity.** Dually labeled *D. discoideum* Ax2 or Δ*gnbp* producing cytosolic mCherry (pDM1042) and P4C-GFP (pWS034) was left untreated or treated with LAI-1 (10 µM, 1 h), or DMSO (solvent control), infected (MOI 5, 4 h) with mCerulean-producing *L. pneumophila* JR32 (pNP99) and analyzed by confocal microscopy. Scale bars, 3 μm. Single channels and merge are shown.

**Fig. S9. Synthesis and application of azido-LAI-1.** (**A**) Reagents and conditions: a) HNMe(OMe)•HCl (1.15 eq.), NMM (1.15 eq.), EDC•HCl (1.15 eq.), CH_2_Cl_2_ –15 °C→rt, 65 h, quant.; b) TBDPSCl (1.15 eq.), imidazole (4.60 eq.), DMF, 0 °C→rt→55 °C, 85%; c) DHP (1.50 eq.), P*p*Ts (0.10 eq.), CH_2_Cl_2_, 0 °C→rt, 19 h, 99%; d), Mg (8.00 eq.), **4** (2.10 eq.), THF, 0 °C→rt, 16 h, 56%; e) P*p*Ts (0.30 eq.), THF:MeOH (3:1), 60 °C, 20 h, 92%; f) CBr_4_ (1.50 eq.), PPh_3_ (1.50 eq.), CH_2_Cl_2_, 0 °C→rt, 17 h, 99%; NaN_3_ (3.00 eq.), DMF, 60 °C, 16 h, quant.; h) TBAF (1 M in THF, 1.20 eq.), THF, 0 °C→rt, 1.5 h, 90%. DHP = 3,4-Dihydro-2*H*-pyran, DMF = *N*,*N*-Dimethylformamide, EDC = 1-Ethyl-3-(3-dimethylaminopropyl)carbodiimide, eq. = equivalents, NMM = 4-Methylmorpholine, P*p*Ts = pyridinium *p*-toluenesulfonate, quant. = quantitative; TBAF = tetra-*n*-butylammonium fluoride; TBDPSCl = *tert-*butyldiphenylsilyl chloride, THF = tetrahydrofuran. (**B**) Azido-LAI-1 can be attached to various conjugation partners (e.g., dyes) using SPAAC.

**Fig. S10. Synthesis and application of diazirine-(*S*)-LAI-1.** (**A**) Reagents and conditions: a) DHP (1.50 eq.), P*p*Ts (0.10 eq.), CH_2_Cl_2_, 0 °C→rt, 18 h, 95%; b) Mg (8.00 eq.), **10** (2.10 eq.), THF, 0 °C→rt, 16 h, 91%; c) P*p*Ts (0.25 eq.), THF:MeOH (3:1), 60 °C, 20 h, 91%; d) CSA (1.00 eq.), 1,2-ethanediol, (24.6 eq.), ethyl orthoformate (8.30 eq.), 50 °C, 16 h, 74%; e) PDC (3.50 eq.), DMF, rt, 16 h, quant.; f) HNMe(OMe)•HCl (1.15 eq.), NMM (1.15 eq.), EDC•HCl (1.15 eq.), CH_2_Cl_2_ 0 °C→rt, 49 h, 72%; g) DHP (1.50 eq.), P*p*Ts (0.10 eq.), CH_2_Cl_2_, 0 °C→rt, 18 h, 87%; h) Mg (8.00 eq.), **16** (2.35 eq.), THF, 0 °C→rt, 17 h, 91%; i) NH_3_ (7 M in MeOH, 75 eq.), HOSA (1.15 eq), MeOH, 0 °C→rt, 51 h; j) I_2_ (1.25 eq.), NEt_3_ (2.00 eq.), MeOH, 0 °C→rt, 16 h, 42% over two steps; k) I_2_ (0.10 eq.), acetone (58.4 eq.), 60 °C, 10 min; l)) P*p*Ts (0.25 eq.), THF:MeOH (3:1), 60 °C, 55 h, 89% over two steps; m) CBr_4_ (1.50 eq.), PPh_3_ (1.50 eq.), CH_2_Cl_2_, 0 °C→rt, 17 h, 73%; n) NaN_3_ (3.00 eq.), DMF, 60 °C, 16 h, 99%; o) TBAF (1 M in THF, 1.20 eq.), THF, 0 °C→rt, 1.5 h, 85%. CSA = camphorsulfonic acid, PDC = pyridinium dichromate, HOSA = hydroxylamine-*O*-sulfonic acid. (**B**) The diazirine function of azido-diazirine-LAI-1 is stimulated by UV light and forms a carbene by releasing nitrogen. The highly reactive carbene can interact with various chemical moieties and thus covalently binds to its biological environment. The covalently fixed azido-LAI-1-derivative can then be attached to various conjugation partners (e.g., dyes, biotin) using SPAAC.

## References

Andreu N, Zelmer A, Fletcher T, Elkington PT, Ward TH, Ripoll J, Parish T, Bancroft GJ, Schaible U, Robertson BD et al (2010) Optimisation of bioluminescent reporters for use with mycobacteria. PLoS One 5: e10777

Asrat S, de Jesus DA, Hempstead AD, Ramabhadran V, Isberg RR (2014) Bacterial pathogen manipulation of host membrane trafficking. Ann Rev Cell Dev Biol 30: 79–109

Barisch C, Paschke P, Hagedorn M, Maniak M, Soldati T (2015) Lipid droplet dynamics at early stages of *Mycobacterium marinum* infection in *Dictyostelium*. Cell Microbiol 17: 1332–1349

Bärlocher K, Hutter CAJ, Swart AL, Steiner B, Welin A, Hohl M, Letourneur F, Seeger MA, Hilbi H (2017) Structural insights into Legionella RidL-Vps29 retromer subunit interaction reveal displacement of the regulator TBC1D5. Nat Commun 8: 1543

Bass AR, Egan MS, Alexander-Floyd J, Lopes Fischer N, Doerner J, Shin S (2023) Human GBP1 facilitates the rupture of the *Legionella*-containing vacuole and inflammasome activation. mBio 14: e0170723

Bloomfield G, Tanaka Y, Skelton J, Ivens A, Kay RR (2008) Widespread duplications in the genomes of laboratory stocks of *Dictyostelium discoideum*. Genome Biol 9: R75

Boamah DK, Zhou G, Ensminger AW, O’Connor TJ (2017) From many hosts, one accidental pathogen: the diverse protozoan hosts of *Legionella*. Front Cell Infect Microbiol 7: 477

Carroll P, Schreuder LJ, Muwanguzi-Karugaba J, Wiles S, Robertson BD, Ripoll J, Ward TH, Bancroft GJ, Schaible UE, Parish T (2010) Sensitive detection of gene expression in mycobacteria under replicating and non-replicating conditions using optimized far-red reporters. PLoS One 5: e9823

Corey EJ, Schmidt G (1979) Useful procedures for the oxidation of alcohols involving pyridinium dichromate in aprotic media. Tetrahedron Lett 20: 399–402

Cosma CL, Humbert O, Ramakrishnan L (2004) Superinfecting mycobacteria home to established tuberculous granulomas. Nat Immunol 5: 828–835

Dunn JD, Bosmani C, Barisch C, Raykov L, Lefrancois LH, Cardenal-Munoz E, Lopez-Jimenez AT, Soldati T (2017) Eat prey, live: *Dictyostelium discoideum* as a model for cell-autonomous defenses. Front Immunol 8: 1906

Fan M, Kiefer P, Charki P, Hedberg C, Seibel J, Vorholt JA, Hilbi H (2023) The *Legionella* autoinducer LAI-1 is delivered by outer membrane vesicles to promote inter-bacterial and inter-kingdom signaling. J Biol Chem 299: 105376

Feeley JC, Gibson RJ, Gorman GW, Langford NC, Rasheed JK, Mackel DC, Baine WB (1979) Charcoal-yeast extract agar: primary isolation medium for *Legionella pneumophila*. J Clin Microbiol 10: 437–441

Fink J, Schumacher F, Schlegel J, Stenzel P, Wigger D, Sauer M, Kleuser B, Seibel J (2021) Azidosphinganine enables metabolic labeling and detection of sphingolipid de novo synthesis. Org Biomol Chem 19: 2203–2212

Goody RS, Itzen A (2013) Modulation of small GTPases by Legionella. Curr Top Microbiol Immunol 376: 117–133

Hilbi H, Buchrieser C (2022) Microbe profile: *Legionella pneumophila* - a copycat eukaryote. Microbiology (Reading) 168: doi: 10.1099/mic.1090.001142

Hilbi H, Hoffmann C, Harrison CF (2011) *Legionella* spp. outdoors: colonization, communication and persistence. Environ Microbiol Rep 3: 286–296

Hochstrasser R, Hilbi H (2017) Intra-species and inter-kingdom signaling of *Legionella pneumophila*. Front Microbiol 8: 79

Hochstrasser R, Hilbi H (2020) *Legionella* quorum sensing meets cyclic-di-GMP signaling. Curr Opin Microbiol 55: 9–16

Hochstrasser R, Hilbi H (2022) The *Legionella* Lqs-LvbR regulatory network controls temperature-dependent growth onset and bacterial cell density. Appl Environ Microbiol 88: e0237021

Hochstrasser R, Hutter CAJ, Arnold FM, Bärlocher K, Seeger MA, Hilbi H (2020) The structure of the *Legionella* response regulator LqsR reveals amino acids critical for phosphorylation and dimerization. Mol Microbiol 113: 1070–1084

Hochstrasser R, Kessler A, Sahr T, Simon S, Schell U, Gomez-Valero L, Buchrieser C, Hilbi H (2019) The pleiotropic *Legionella* transcription factor LvbR links the Lqs and c-di-GMP regulatory networks to control biofilm architecture and virulence. Environ Microbiol 21: 1035–1053

Hochstrasser R, Michaelis S, Brülisauer S, Sura T, Fan M, Maass S, Becher D, Hilbi H (2022) Migration of *Acanthamoeba* through *Legionella* biofilms is regulated by the bacterial Lqs-LvbR network, effector proteins and the flagellum. Environ Microbiol 24: 3672–3692

Hoffmann C, Finsel I, Otto A, Pfaffinger G, Rothmeier E, Hecker M, Becher D, Hilbi H (2014a) Functional analysis of novel Rab GTPases identified in the proteome of purified *Legionella*-containing vacuoles from macrophages. Cell Microbiol 16: 1034–1052

Hoffmann C, Harrison CF, Hilbi H (2014b) The natural alternative: protozoa as cellular models for Legionella infection. Cell Microbiol 16: 15–26

Horwitz MA (1983) Formation of a novel phagosome by the Legionnaires’ disease bacterium (*Legionella pneumophila*) in human monocytes. J Exp Med 158: 1319–1331.

Hubber A, Roy CR (2010) Modulation of host cell function by *Legionella pneumophila* type IV effectors. Ann Rev Cell Dev Biol 26: 261–283

Hüsler D, Stauffer P, Hilbi H (2023a) Tapping lipid droplets: A rich fat diet of intracellular bacterial pathogens. Mol Microbiol 120: 194–209

Hüsler D, Stauffer P, Keller B, Böck D, Steiner T, Ostrzinski A, Vormittag S, Striednig B, Swart AL, Letourneur F et al (2023b) The large GTPase Sey1/atlastin mediates lipid droplet- and FadL-dependent intracellular fatty acid metabolism of *Legionella pneumophila*. eLife 12: e85142

Hüsler D, Steiner B, Welin A, Striednig B, Swart AL, Molle V, Hilbi H, Letourneur F (2021) *Dictyostelium* lacking the single atlastin homolog Sey1 shows aberrant ER architecture, proteolytic processes and expansion of the *Legionella*-containing vacuole. Cell Microbiol 23: e13318

Journet A, Klein G, Brugiere S, Vandenbrouck Y, Chapel A, Kieffer S, Bruley C, Masselon C, Aubry L (2012) Investigating the macropinocytic proteome of *Dictyostelium* amoebae by high-resolution mass spectrometry. Proteomics 12: 241–245

Kagan JC, Roy CR (2002) *Legionella* phagosomes intercept vesicular traffic from endoplasmic reticulum exit sites. Nat Cell Biol 4: 945–954

Katic A, Hüsler D, Letourneur F, Hilbi H (2021) *Dictyostelium* dynamin superfamily GTPases implicated in vesicle trafficking and host-pathogen interactions. Front Cell Dev Biol 9: 731964

Kendall MM, Sperandio V (2016) What a dinner party! Mechanisms and functions of interkingdom signaling in host-pathogen associations. mBio 7: e01748

Kessler A, Schell U, Sahr T, Tiaden A, Harrison C, Buchrieser C, Hilbi H (2013) The *Legionella pneumophila* orphan sensor kinase LqsT regulates competence and pathogen-host interactions as a component of the LAI-1 circuit. Environ Microbiol 15: 646–662

Kirkby M, Enosi Tuipulotu D, Feng S, Lo Pilato J, Man SM (2023) Guanylate-binding proteins: mechanisms of pattern recognition and antimicrobial functions. Trends Biochem Sci 48: 883–893

Knobloch P, Koliwer-Brandl H, Arnold FM, Hanna N, Gonda I, Adenau S, Personnic N, Barisch C, Seeger MA, Soldati T et al (2020) *Mycobacterium marinum* produces distinct mycobactin and carboxymycobactin siderophores to promote growth in broth and phagocytes. Cell Microbiol 22: e13163

Koliwer-Brandl H, Knobloch P, Barisch C, Welin A, Hanna N, Soldati T, Hilbi H (2019) Distinct *Mycobacterium marinum* phosphatases determine pathogen vacuole phosphoinositide pattern, phagosome maturation, and escape to the cytosol. Cell Microbiol 21: e13008

Kutsch M, Coers J (2021) Human guanylate binding proteins: nanomachines orchestrating host defense. FEBS J 288: 5826–5849

Kutsch M, Sistemich L, Lesser CF, Goldberg MB, Herrmann C, Coers J (2020) Direct binding of polymeric GBP1 to LPS disrupts bacterial cell envelope functions. EMBO J 39: e104926

Laevsky G, Knecht DA (2001) Under-agarose folate chemotaxis of *Dictyostelium discoideum* amoebae in permissive and mechanically inhibited conditions. Biotechniques 31: 1140–1142, 1144, 1146-1149

Lang J, Bohn P, Bhat H, Jastrow H, Walkenfort B, Cansiz F, Fink J, Bauer M, Olszewski D, Ramos-Nascimento A et al (2020) Acid ceramidase of macrophages traps herpes simplex virus in multivesicular bodies and protects from severe disease. Nat Commun 11: 1338

Liu BC, Sarhan J, Panda A, Muendlein HI, Ilyukha V, Coers J, Yamamoto M, Isberg RR, Poltorak A (2018) Constitutive interferon maintains GBP expression required for release of bacterial components upstream of pyroptosis and anti-DNA responses. Cell Rep 24: 155–168 e155

Lockwood DC, Amin H, Costa TRD, Schroeder GN (2022) The *Legionella pneumophila* Dot/Icm type IV secretion system and its effectors. Microbiology (Reading) 168: doi: 10.1099/mic.1090.001187

Loovers HM, Kortholt A, de Groote H, Whitty L, Nussbaum RL, van Haastert PJ (2007) Regulation of phagocytosis in *Dictyostelium* by the inositol 5-phosphatase OCRL homolog Dd5P4. Traffic 8: 618–628

Lu H, Clarke M (2005) Dynamic properties of *Legionella*-containing phagosomes in *Dictyostelium* amoebae. Cell Microbiol 7: 995–1007

Manstein DJ, Schuster HP, Morandini P, Hunt DM (1995) Cloning vectors for the production of proteins in *Dictyostelium discoideum*. Gene 162: 129–134

Michaelis S, Chen T, Schmid C, Hilbi H (2024a) Nitric oxide signaling through three receptors regulates virulence, biofilm formation, and phenotypic heterogeneity of *Legionella pneumophila*. mBio 15: e0071024

Michaelis S, Gomez-Valero L, Chen T, Schmid C, Buchrieser C, Hilbi H (2024b) Small molecule communication of *Legionella*: the ins and outs of autoinducer and nitric oxide signaling. Microb Mol Biol Rev: e0009723. doi: 10.1128/mmbr.00097-23

Miller MB, Skorupski K, Lenz DH, Taylor RK, Bassler BL (2002) Parallel quorum sensing systems converge to regulate virulence in *Vibrio cholerae*. Cell 110: 303–314

Mondino S, Schmidt S, Rolando M, Escoll P, Gomez-Valero L, Buchrieser C (2020) Legionnaires’ disease: state of the art knowledge of pathogenesis mechanisms of *Legionella*. Ann Rev Pathol 15: 439–466

Nagai H, Kagan JC, Zhu X, Kahn RA, Roy CR (2002) A bacterial guanine nucleotide exchange factor activates ARF on *Legionella phagosomes*. Science 295: 679–682

Newton HJ, Ang DK, van Driel IR, Hartland EL (2010) Molecular pathogenesis of infections caused by *Legionella pneumophila*. Clin Microbiol Rev 23: 274–298

Ng WL, Bassler BL (2009) Bacterial quorum-sensing network architectures. Ann Rev Genet 43: 197–222

Pacheco AR, Sperandio V (2009) Inter-kingdom signaling: chemical language between bacteria and host. Curr Opin Microbiol 12: 192–198

Perry CJ, Warren EC, Damstra-Oddy JL, Storey C, Francione LM, Annesley SJ, Fisher PR, Muller-Taubenberger A, Williams RSB (2020) A *Dictyostelium discoideum* mitochondrial fluorescent tagging vector that does not affect respiratory function. Biochem Biophys Rep 22: 100751

Personnic N, Bärlocher K, Finsel I, Hilbi H (2016) Subversion of retrograde trafficking by translocated pathogen effectors. Trends Microbiol 24: 450–462

Personnic N, Striednig B, Hilbi H (2018) *Legionella* quorum sensing and its role in pathogen-host interactions. Curr Opin Microbiol 41: 29–35

Personnic N, Striednig B, Hilbi H (2021) Quorum sensing controls persistence, resuscitation, and virulence of *Legionella* subpopulations in biofilms. ISME J 15: 196–210

Personnic N, Striednig B, Lezan E, Manske C, Welin A, Schmidt A, Hilbi H (2019) Quorum sensing modulates the formation of virulent *Legionella* persisters within infected cells. Nat Commun 10: 5216

Procacci B, Roy SS, Norcott P, Turner N, Duckett SB (2018) Unlocking a diazirine long-lived nuclear singlet state via photochemistry: NMR detection and lifetime of an unstabilized diazo-compound. J Am Chem Soc 140: 16855–16864

Qiu J, Luo ZQ (2017) *Legionella* and *Coxiella* effectors: strength in diversity and activity. Nat Rev Microbiol 15: 591–605

Rafeld HL, Kolanus W, van Driel IR, Hartland EL (2021) Interferon-induced GTPases orchestrate host cell-autonomous defence against bacterial pathogens. Biochem Soc Trans 49: 1287–1297

Ragaz C, Pietsch H, Urwyler S, Tiaden A, Weber SS, Hilbi H (2008) The *Legionella pneumophila* phosphatidylinositol-4 phosphate-binding type IV substrate SidC recruits endoplasmic reticulum vesicles to a replication-permissive vacuole. Cell Microbiol 10: 2416–2433

Robinson CG, Roy CR (2006) Attachment and fusion of endoplasmic reticulum with vacuoles containing *Legionella pneumophila*. Cell Microbiol 8: 793–805

Rühling M, Kersting L, Wagner F, Schumacher F, Wigger D, Helmerich DA, Pfeuffer T, Elflein R, Kappe C, Sauer M et al (2024) Trifunctional sphingomyelin derivatives enable nanoscale resolution of sphingomyelin turnover in physiological and infection processes via expansion microscopy. Nat Commun 15: 7456

Sadosky AB, Wiater LA, Shuman HA (1993) Identification of *Legionella pneumophila* genes required for growth within and killing of human macrophages. Infect Immun 61: 5361–5373

Santos JC, Boucher D, Schneider LK, Demarco B, Dilucca M, Shkarina K, Heilig R, Chen KW, Lim RYH, Broz P (2020) Human GBP1 binds LPS to initiate assembly of a caspase-4 activating platform on cytosolic bacteria. Nat Commun 11: 3276

Schell U, Kessler A, Hilbi H (2014) Phosphorylation signalling through the *Legionella* quorum sensing histidine kinases LqsS and LqsT converges on the response regulator LqsR. Mol Microbiol 92: 1039–1055

Schell U, Simon S, Sahr T, Hager D, Albers MF, Kessler A, Fahrnbauer F, Trauner D, Hedberg C, Buchrieser C et al (2016) The alpha-hydroxyketone LAI-1 regulates motility, Lqs-dependent phosphorylation signalling and gene expression of *Legionella pneumophila*. Mol Microbiol 99: 778–793

Segal G, Shuman HA (1998) Intracellular multiplication and human macrophage killing by *Legionella pneumophila* are inhibited by conjugal components of IncQ plasmid RSF1010. Mol Microbiol 30: 197–208

Shank EA, Kolter R (2009) New developments in microbial interspecies signaling. Curr Opin Microbiol 12: 205–214

Simon S, Schell U, Heuer N, Hager D, Albers MF, Matthias J, Fahrnbauer F, Trauner D, Eichinger L, Hedberg C et al (2015) Inter-kingdom signaling by the *Legionella* quorum sensing molecule LAI-1 modulates cell migration through an IQGAP1-Cdc42-ARHGEF9-dependent pathway. PLoS Pathog 11: e1005307

Simon S, Wagner MA, Rothmeier E, Müller-Taubenberger A, Hilbi H (2014) Icm/Dot-dependent inhibition of phagocyte migration by *Legionella* is antagonized by a translocated Ran GTPase activator. Cell Microbiol 16: 977–992

Spirig T, Tiaden A, Kiefer P, Buchrieser C, Vorholt JA, Hilbi H (2008) The *Legionella* autoinducer synthase LqsA produces an α-hydroxyketone signaling molecule. J Biol Chem 283: 18113–18123

Steiner B, Swart AL, Welin A, Weber S, Personnic N, Kaech A, Freyre C, Ziegler U, Klemm RW, Hilbi H (2017) ER remodeling by the large GTPase atlastin promotes vacuolar growth of *Legionella pneumophila*. EMBO Rep 18: 1817–1836

Steiner B, Weber S, Hilbi H (2018) Formation of the *Legionella*-containing vacuole: phosphoinositide conversion, GTPase modulation and ER dynamics. Int J Med Microbiol 308: 49–57

Sternstein C, Böhm TM, Fink J, Meyr J, Ludemann M, Krug M, Kriukov K, Gurdap CO, Sezgin E, Ebert R et al (2023) Development of an effective functional lipid anchor for membranes (FLAME) for the bioorthogonal modification of the lipid bilayer of mesenchymal stromal cells. Bioconjug Chem 34: 1221–1233

Striednig B, Hilbi H (2022) Bacterial quorum sensing and phenotypic heterogeneity: how the collective shapes the individual. Trends Microbiol 30: 379–389

Striednig B, Lanner U, Niggli S, Katic A, Vormittag S, Brülisauer S, Hochstrasser R, Kaech A, Welin A, Flieger A et al (2021) Quorum sensing governs a transmissive *Legionella* subpopulation at the pathogen vacuole periphery. EMBO Rep 22: e52972

Sun J, Dong Y, Cao L, Wang X, Wang S, Hu Y (2004) Highly efficient chemoselective deprotection of O,O-acetals and O,O-ketals catalyzed by molecular iodine in acetone. J Org Chem 69: 8932–8934

Swart AL, Gomez-Valero L, Buchrieser C, Hilbi H (2020) Evolution and function of bacterial RCC1 repeat effectors. Cell Microbiol 22: e13246

Swart AL, Harrison CF, Eichinger L, Steinert M, Hilbi H (2018) *Acanthamoeba* and *Dictyostelium* as cellular models for *Legionella* infection. Front Cell Infect Microbiol 8: 61

Swart AL, Hilbi H (2020) Phosphoinositides and the fate of *Legionella* in phagocytes. Front Immunol 11: 25

Tiaden A, Hilbi H (2012) α-Hydroxyketone synthesis and sensing by *Legionella* and *Vibrio*. Sensors 12: 2899–2919

Tiaden A, Spirig T, Carranza P, Brüggemann H, Riedel K, Eberl L, Buchrieser C, Hilbi H (2008) Synergistic contribution of the *Legionella pneumophila lqs* genes to pathogen-host interactions. J Bacteriol 190: 7532–7547

Tiaden A, Spirig T, Hilbi H (2010a) Bacterial gene regulation by α-hydroxyketone signaling. Trends Microbiol 18: 288–297

Tiaden A, Spirig T, Sahr T, Wälti MA, Boucke K, Buchrieser C, Hilbi H (2010b) The autoinducer synthase LqsA and putative sensor kinase LqsS regulate phagocyte interactions, extracellular filaments and a genomic island of *Legionella pneumophila*. Environ Microbiol 12: 1243–1259

Tiaden A, Spirig T, Weber SS, Brüggemann H, Bosshard R, Buchrieser C, Hilbi H (2007) The *Legionella pneumophila* response regulator LqsR promotes host cell interactions as an element of the virulence regulatory network controlled by RpoS and LetA. Cell Microbiol 9: 2903–2920

Tretina K, Park ES, Maminska A, MacMicking JD (2019) Interferon-induced guanylate-binding proteins: guardians of host defense in health and disease. J Exp Med 216: 482–500

Veltman DM, Akar G, Bosgraaf L, Van Haastert PJM (2009) A new set of small, extrachromosomal expression vectors for *Dictyostelium discoideum*. Plasmid 61: 110–118

Vormittag S, Ende RJ, Derré I, Hilbi H (2023a) Pathogen vacuole membrane contact sites - close encounters of the fifth kind. Microlife 4: uqad018

Vormittag S, Hüsler D, Haneburger I, Kroniger T, Anand A, Prantl M, Barisch C, Maass S, Becher D, Letourneur F et al (2023b) *Legionella*- and host-driven lipid flux at LCV-ER membrane contact sites promotes vacuole remodeling. EMBO Rep 24: e56007

Wang L, Ishida A, Hashidoko Y, Hashimoto M (2017) Dehydrogenation of the NH-NH bond triggered by potassium tert-butoxide in liquid ammonia. Angew Chem Int Ed Engl 56: 870–873

Weber S, Hilbi H (2014) Live cell imaging of phosphoinositide dynamics during *Legionella* infection. Methods Mol Biol 1197: 153–167

Weber S, Steiner B, Welin A, Hilbi H (2018) *Legionella*-containing vacuoles capture PtdIns(4)*P*-rich vesicles derived from the Golgi apparatus. mBio 9: e02420–18

Weber S, Wagner M, Hilbi H (2014) Live-cell imaging of phosphoinositide dynamics and membrane architecture during *Legionella* infection. mBio 5: e00839–13

Weber SS, Ragaz C, Hilbi H (2009) The inositol polyphosphate 5-phosphatase OCRL1 restricts intracellular growth of *Legionella*, localizes to the replicative vacuole and binds to the bacterial effector LpnE. Cell Microbiol 11: 442–460

Weber SS, Ragaz C, Reus K, Nyfeler Y, Hilbi H (2006) *Legionella pneumophila* exploits PI(4)*P* to anchor secreted effector proteins to the replicative vacuole. PLoS Pathog 2: e46

Welin A, Weber S, Hilbi H (2018) Quantitative imaging flow cytometry of *Legionella*-infected *Dictyostelium* amoebae reveals the impact of retrograde trafficking on pathogen vacuole composition. Appl Environ Microbiol 84: e00158–18

Wiegand S, Kruse J, Gronemann S, Hammann C (2011) Efficient generation of gene knockout plasmids for *Dictyostelium discoideum* using one-step cloning. Genomics 97: 321–325

Yang Z, Attygalle AB, Meinwald J (2000) Reptilian chemistry: enantioselective syntheses of novel components from a crocodile exocrine secretion. Synthesis 2000: 1936–1943

